# *Fasciola hepatica* Fatty Acid Binding Protein 1 modulates T cell polarization by promoting dendritic cell thrombospondin-1 secretion without affecting metabolic homeostasis in obese mice

**DOI:** 10.1101/2022.04.01.486671

**Authors:** Anna Zawistowska-Deniziak, Joost M. Lambooij, Alicja Kalinowska, Thiago A. Patente, Maciej Łapiński, Hendrik J.P. van der Zande, Katarzyna Basałaj, Clarize de Korne, Mathilde A.M. Chayé, Tom Gasan, Luke J. Norbury, Martin Giera, Arnaud Zaldumbide, Hermelijn H. Smits, Bruno Guigas

## Abstract

**Background:** The parasitic trematode *Fasciola hepatica* evades host immune defenses through secretion of various immunomodulatory molecules. Fatty Acid Binding Proteins (*fh*FABPs) are among the main excreted/secreted proteins and have been shown to display anti-inflammatory properties. However, little is currently known regarding their impact on dendritic cells (DCs) and their subsequent capacity to prime specific CD4^+^ T cell subsets.

**Methodology/Principal Findings:** The immunomodulatory effects of both native *F. hepatica* extracts and recombinant *fh*FABPs were assessed on monocyte-derived human DCs (moDCs) and the underlying mechanism was next investigated using various approaches, including DC-allogenic T cell co-culture and DC phenotyping through transcriptomic, proteomic and FACS analyses. We mainly showed that *fh*FABP1 induced a tolerogenic-like phenotype in LPS-stimulated moDCs characterized by a dose-dependent increase in the cell-surface tolerogenic marker CD103 and IL-10 secretion, while DC co-stimulatory markers were not affected. A significant decrease in secretion of the pro-inflammatory cytokines IL-12p70 and IL-6 was also observed. In addition, these effects were associated with an increase in both Th2-on-Th1 ratio and IL-10 secretion by CD4^+^ T cells following DC-T cell co-culture. RNA sequencing and targeted proteomic analyses identified thrombospondin-1 (TSP-1) as a non-canonical factor highly expressed and secreted by *fh*FABP1-primed moDCs. The effect of *fh*FABP1 on T cell skewing was abolished when using a TSP-1 blocking antibody during DC-T cell co-culture. Immunomodulation by helminth molecules has been linked to improved metabolic homeostasis during obesity. Although *fh*FABP1 injection in high-fat diet-fed obese mice induced a potent Th2 immune response in adipose tissue, it did not improved insulin sensitivity or glucose homeostasis.

**Conclusions/Significance:** We show that *fh*FABP1 modulates T cell polarization, notably by promoting DC TSP-1 secretion *in vitro*, without affecting metabolic homeostasis in a mouse model of type 2 diabetes.

## Introduction

Helminths are master regulators of the immune system, allowing them to infect millions of humans and other animals worldwide [1]. Manipulation of the host immune system by the parasites occurs at different levels through various mechanisms developed during coevolution [2; 3]. Usually, the helminth-induced immune responses allows maintenance of long-term infection, striking a balance between parasite feeding and reproduction and minimal damage to the host. The classical immune response to helminth infections is the induction of type 2 immunity, characterized by high eosinophilia and production of IL-4, IL-5 and IL-13, together with a network of IL-10-producing regulatory B and T cells that both contribute to the prevention of excessive inflammation and tissue damage inflicted by the worms [4; 5]. We and others have also reported that various helminth molecules can trigger a potent type 2 immune response in metabolic organs and improve insulin sensitivity and glucose homeostasis in metabolically-impaired conditions [6]. Professional antigen presenting cells (APCs), especially dendritic cells (DCs), are central players in initiating and regulating these specific adaptive immune responses [7; 8; 9]. Helminth-derived molecules, especially those present in excretory-secretory products (ESP), are thought to interact with DCs, notably via both pattern recognition receptor (PRR)-dependent and independent mechanisms, and modulate DCs capacity to prime different CD4^+^ T cell subsets [5]. Although comprehensive omics analyses have helped to identify and characterize the molecules present in the tissue extracts and ESP from various helminth species [10; 11; 12; 13], there is still a significant lack in knowledge regarding which and how specific helminth-derived molecules modulate DC-mediated immune responses.

One of the most prevalent helminths with zoonotic potential is *Fasciola hepatica*, which currently affects more than 17 million people worldwide, with an additional 180 million humans being at risk of infection, consequently, its associated disease fascioliasis [14]. The early stages of infection are associated with a mixed Th1/Th2 response characterized by increased IFNγ, IL-4, IL-10 and TGF-β-producing CD4^+^ T cells. When the infection progresses the type 2 immune response becomes predominant which coincides with suppression of Th1-induced inflammation and an induction of regulatory T cells (Treg) [15; 16; 17]. Interestingly, it has been reported that both *F. hepatica* tegumental antigens and ESPs (*fh*ESPs) can suppress the maturation of DCs [18; 19; 20; 21], and induce T cell anergy [22; 23]. Additionally, the glycans present on *F. hepatica-derived* molecules have been shown to play a role in driving the Th2/Treg immune responses by interacting with CD209 (DC-SIGN) and CD206 (MR) C-type lectin receptors on APCs [15; 22; 24; 25; 26]. Proteomic analyses of *fh*ESPs identified Fatty Acid Binding Proteins (*fh*FABPs) as some of the main excreted/secreted proteins, together with cathepsin Ls and antioxidant molecules (*fh*GST and *fh*TPX) [27; 28; 29]. Interestingly, *fh*FABPs have been shown to display antiinflammatory properties, notably in macrophages [30]. However, little is currently known regarding their impact on human DCs and their subsequent capacity to prime specific CD4+ T cell subsets.

In the present study, we mainly assessed the immunomodulatory effects of yeast-produced recombinant *F. hepatica* molecules, especially single FABP isoforms, on monocyte-derived human DCs (moDCs). Subsequently, we investigated the underlying mechanism of *fh*FABP1 using a combination of experimental approaches, including in depth DC phenotyping through transcriptomic, proteomic and FACS analyses. Our data show that *fh*FABP1 induces a tolerogenic-like phenotype in human moDCs and modulates T cell polarization by promoting DC thrombospondin-1 (TSP-1) secretion. However, despite inducing a Th2 cell response *in vivo*, fhFABP1 did not affect metabolic homeostasis in obese mice, a model of insulin resistance and type 2 diabetes.

## Materials and Methods

### Preparation and purification of *F. hepatica* derived molecules

Adult *F. hepatica* flukes were collected from the liver of infected sheep, as previously described [31]. The whole *F. hepatica* tissue extract containing a mixture of somatic antigens (FhSA) was obtained by homogenizing two flukes in sterile PBS using a glass homogenizer, followed by a sonification step (4 x 30 sec with intervals of 20 sec on ice) and centrifugation for 25 min (18 000 x *g* at 4°C). *fh*ESPs were collected as described previously [31]. Briefly, adult flukes were first washed in PBS and incubated in RPMI 1640 at 37°C for 2 hours, then incubated for ^~^32 hours at 37°C in RPMI 1640 supplemented with 100 U/ml of penicillin and 0.1 mg/mlof streptomycin. The media was collected and immediately frozen at −80°C, and replaced with fresh media every 2-3 hours. Later, batches were combined, spun down (8,000 × *g*, 10 min, 4°C) and concentrated until ^~^1.5 mg/ml using Amicon Ultra-15 Centrifugal Filter Unit with a 3 kDa cut-off (Millipore). Protein concentration was determined by BCA assay (Pierce/Thermo Scientific).

### FABP cloning and recombinant protein production

Total RNA was isolated from adult *F. hepatica* fluke using a Total RNA mini Plus kit (A&A Biotechnology) and reverse transcribed into cDNAs using a Maxima^™^ First Strand cDNA Synthesis Kit for RT-qPCR (Thermo Scientific) according to the manufacturer’s protocol. The total cDNA was used as a template for FABP1 (GenBank: M95291.1) and FABP5 (GenBank: KJ713302.1) amplification, respectively. Full coding sequences were cloned in frame into the pPICZαB vector (Invitrogen) using PstI and XbaI restriction sites. The primer sequences used were: [F_FhFABP1_PstI] ATT CTG CAG GGA TGG CTG ACT TTG TGG, [R_FhFABP1_XbaI] TCT CTA GAC TCG CTT TGA GCA GAG TGG, [F_FhFABP5_PstI] CCG CTG CAG TCA TGT CTG GAT TTA TCG G, [R_FhFABP5_XbaI] GGC TCT AGA TCT TTG ATG CGT TGG TAT CTC C. The integrity of the constructs was verified by nucleotide sequencing. Recombinant plasmids were then linearized with PmeI and transformed into *Pichia pastoris* X33 strain using electroporation method. The expression of recombinant FABPs was performed in Buffered complex medium containing 0.5% methanol at 29°C for 96 h. The protein was purified on a Ni-NTA resin column (Macherey-Nagel) and the eluted fractions were concentrated and dialyzed against PBS using the same Amicon system as described above. Endotoxins were removed using Endotoxin Removal Spin Columns (Pierce) and the final eluate was passed through a 0.22 μm filter. The presence of purified recombinant FABPs was confirmed by SDS-PAGE and Western blotting analysis. Cathepsin L5 (*fh*CL5) was produced as previously described [32]. Protein concentrations were determined by BCA assay (Pierce/Thermo).

### Human moDC culture, stimulation and analysis

Human monocytes were isolated from peripheral venous blood/buffy coats purchased from Sanquin (The Netherlands) by gradient centrifugation and positive selection using CD14 MACS beads (Miltenyi Biotec, Germany), as previously described [33]. Monocytes were next differentiated to moDCs using human rGM-CSF (20ng/ml, BioCource/Life technologies) and rIL-4 (0.86ng/ml, R&D Systems) for 5-6 days, with media refreshment on day 3. Immature moDC (0.5×10^6^/ml) were stimulated with 100 ng/ml LPS (*E. coli* 0111 B4 strain, InvivoGen, San Diego) on day 5-6 in the presence or absence of *F. hepatica* molecules *fh*SA (50 μg/ml), *fh*ESPs (25 μg/ml), recombinant molecules *fh*FABP1 (10, 25, 50 μg/ml), *fh*FABP5 and *fh*CL5 (25 μg/ml each). After 48 hours of stimulation, cell supernatants were collected for cytokine/chemokine analyses and cells were stained with Fixable Aqua Dead Cell Stain Kit (Invitrogen/Thermo Scientific) for determination of cell surface expression of co-stimulatory molecules by FACS (FACSCanto II, BD Bioscience) using various antibodies **(Table S1).** In addition, primed moDC (1×10^4^ cells) were co-cultured with CD40L-expressing J558 cells (1:1) for 24 hours, followed by supernatant collection. Cytokine (IL-12p70, IL-6, IL-10) and chemokine (CXCL11, TSP-1) concentrations were determined using commercial ELISA kits **(Table S1).**

### Human T cell polarization

For analysis of T cell polarization, moDCs primed with LPS and *F. hepatica* molecules were co-cultured with allogenic naïve CD4+ T cells for 6-7 days in the presence of staphylococcal enterotoxin B (10 pg/ml) with or without the addition of a chemokine mix (CCL19, CXCL9, CXCL11; 0.33ng/ml of each (Peprotech), α-TSP1 (clone 133, Sigma-Aldrich) or isotype control IgG2b (BioLegend, Ultra LEAF Purified) antibodies (10 μg/ml). On day 7, cells were harvested and transferred to a 24 well plate and cultured in the presence of human rIL-2 (10U/ml, R&D Systems) for 4 additional days. Intracellular cytokine production was analyzed after 6 hours restimulation with phorbol myristate acetate (100 ng/ml) and ionomycin (1 μg/ml) with addition of brefeldin A (10 μg/ml) during the last 4 hours. Subsequently, the cells were stained with Fixable Aqua Dead Cell Stain Kit (Invitrogen) for 15 min at RT, fixed with 1.9% formaldehyde (Sigma-Aldrich), permeabilized (eBioscience^™^ Permeabilization Buffer) and stained with anti-IL-4, anti-IL-10 and anti-IFNγ for 30 min at RT. Alternatively, 1×10^5^ T cells were restimulated using anti-CD3 and anti-CD28 (BD Biosciences). 24 hours after restimulation, supernatants were collected, and IL-10 production by T cells was measured by ELISA (BD Bioscience). For analysis of transcription factors, cells were first stained with Fixable Aqua Dead Cell Stain Kit (Invitrogen) for 15 min at RT and fixed with FoxP3/Transcription Factor Staining Buffer Set (Invitrogen, for FOXP3 detection) for 1 hour at 4°C. Then, cells were washed twice with permeabilization buffer (eBioscience) and stained with CD3, CD4, FOXP3, GATA3, and T-bet antibodies **(Table S1).** All samples were run on a FACSCanto II or BD LSR II and analyzed using FlowJo (Version vX.0.7 TreeStar).

### T cell suppression assay

To analyze the suppressive capacity of moDCprimed T cells, 5 x 10^4^ moDCs were co-cultured with 5 x 10^5^ naïve CD4^+^ T cells for 6 days. These T cells (test T cells) were harvested, washed, counted and irradiated (3000 RAD) to prevent expansion. Bystander target T cells (responder T cells), which were allogeneic memory T cells from the same donor as the test T cells, were labeled with 0.5 μM cell tracking dye 5,6-carboxy fluorescein diacetate succinimidyl ester (CFSE). Subsequently, 5 x 10^4^ test T cells, 2.5 x 10^4^ responder T cells, and 1 x 10^3^ LPS-stimulated DC were co-cultured for an additional 6 days. Proliferation was determined by flow cytometry, by co-staining with CD3, CD4 and CD25 antibodies **(Table S1).**

### Cytokine Arrays

The supernatants collected from LPS-stimulated moDCs with or without *fh*FABP1 (25 μg/ml) were pooled from 4 donors in equal volume and subjected to the Proteome Profiler Human XL Cytokine Kit (R&D Systems) according to the manufacturer’s instructions. Pre-blocked nitrocellulose membranes of the Human Cytokine Array were incubated with 1 ml of pooled supernatants and detection antibody cocktail overnight at 4°C on a rocking platform. The membranes were washed three times with 1× Wash Buffer (R&D Systems) to remove unbound proteins. Chemiluminescent detection reagents were applied to detect spot densities. Membranes were exposed to X-ray film and analyzed using the manufacturer’s image analysis software (Quantity One).

### RNA isolation and sequencing and computational analysis

Immature moDC (0.5 x 10^6^/ml) were stimulated with 100 ng/ml LPS on day 5-6 in the presence or absence of *fh*FABP1 (25μg/ml). After 5 h of stimulation, cells were harvested, washed two times with ice-cold PBS and snap-frozen. Total RNA was isolated with an RNA purification kit (Macherey-Nagel) and was subjected to poly(A) enrichment using NEBNext Poly(A) mRNA Magnetic Isolation Module (New England Biolabs). Strand-specific cDNA libraries were constructed with NEBNext^®^ Ultra^™^ II Directional RNA Library Prep Kit for Illumina^®^ (New England Biolabs). Sequencing was performed using NovaSeq6000 (Illumina) sequencer in PE150 mode. The subsequent quality control steps and computational analyses are described in details in the Supplementary method. Data are publicly available on NCBI (GSE197462).

### Animal experiments

All mouse experiments were performed in accordance with the Guide for the Care and Use of Laboratory Animals of the Institute for Laboratory Animal Research and have received approval from the university Ethical Review Board (Leiden University Medical Center, Leiden, The Netherlands; DEC12199 & PE.18.025.010). All mice were purchased from Envigo (Horst, The Netherlands) and housed at Leiden University Medical Center in a temperature-controlled room with a 12 hour light-dark cycle and ad libitum access to food and tap water.

#### Hock injection

8-10 weeks-old C57BL6/J male mice were injected in either left or right hind hock with 30 μl of an emulsion containing 25 μg of *fh*FABP1 or PBS with or without 200 ng of LPS. The draining popliteal lymph nodes (LNs) were collected after 6 days in 500 μl HBSS medium for subsequent analysis. LNs were mashed up with the back of a syringe and incubated with digestion buffer (1 μg/ml of Collagenase D and 2000 U/ml of RNAse) for 20 min at 37°C. Next, samples were filtered through a 100 μm cell strainer and washed 3 times with MACS buffer (PBS, 0.5% BSA, 2 mM EDTA). Cells were ultimately centrifuged at 1500 rpm for 5 min at 4°C, counted and plated for further experiments.

#### Treatment of HFD-induced obese mice

8-10 weeks-old C57BL/6JOlaHsd male mice were fed a high-fat diet (HFD, 45% energy derived from fat, D12451) for 12 weeks. At the end of this run-in period, mice were randomized in two groups based on body weight, fat mass, and fasting plasma glucose levels. *fh*FABP1 (25 μg) or vehicle control (sterile-filtered PBS) and next received intraperitoneal (i.p.) injection every 3 days for 4 weeks. Body weight was monitored throughout the experiment and body composition was measured at week 4 of treatment using an EchoMRI (Echo Medical Systems, Houston, TX, USA). Whole-body insulin tolerance (ipITT) and glucose tolerance (ipGTT) tests were performed at week 3-4 of treatment in 4 hours- or 6 hours-fasted mice, respectively, as described previously [34]. At the end of the experiment, mice were sacrificed through an overdose of ketamine/xylazine and both visceral white adipose tissue (epidydimal; eWAT) and liver were weighed and collected for further processing and analyses.

### *In vitro* restimulation of LN cells

1.5 x 10^6^ popliteal LN cells/ml from individual animals were restimulated with 50 ng/ml PMA plus 2 μg/ml ionomycin and 10 μg/ml brefeldin A for 4 hours. Then cells were first stained with Fixable Aqua Dead Cell Stain Kit (Invitrogen/Thermo) and fixed with 1.9% PFA, and next permeabilized with permeabilization buffer (eBioscience). To analyze CD4^+^ T cells, LN cells were stained for CD11b, CD11c, GR-1, B220, NK1.1 as a lineage cocktail. CD4^+^ T cells were identified as CD3, CD4 and CD44 positive and analyzed for IL-4, IFNγ and IL-10 intracellular expression with specific antibodies **(Table S1).** Alternatively, 1 x 10^6^ popliteal LN cells/ml were stimulated with 10 μg/ml of PHA for 4 days and cytokines levels were measured by FACS using Cytometric Bead Assay (CBA; BD bioscience).

### Immune cell phenotyping in metabolic organs

eWAT and liver were collected at sacrifice after a 1 min transcardial perfusion with PBS through the left ventricle and processed as described previously, with a modified digestion mix for eWAT containing 1 mg/ml collagenase type I from *Clostridium histolyticum*, 2% (w/v) dialyzed bovine serum albumin (BSA, fraction V) and 6 mM D-Glucose in HEPES-buffered Krebs [34; 35]. After isolation, leukocytes were counted using a hemocytometer, incubated with the live/dead marker Zombie-UV, fixed using either 1.85% formaldehyde or the FOXP3 Transcription Factor Staining Buffer kit (Invitrogen) and stained as described below. Cells were measured within 4 days post fixation. For analysis of macrophages, cells were permeabilized and incubated with an antibody against YM1 conjugated to biotin and rabbit-anti-RELMα, washed, and stained with streptavidin-PerCP-Cy5.5, anti-rabbit-Alexa Fluor 647 and antibodies directed against CD45, CD64, F4/80, Ly6C and Siglec-F. For analysis of CD4^+^ T cells, cells were stained with antibodies against CD3, CD4, CD8, CD44, CD45, and a lineage cocktail containing B220, CD11b, CD11c, GR-1 and NK1.1 targeting antibodies. For intracellular cytokine detection, isolated cells were restimulated with 100 ng/ml PMA and 1 μg/ml ionomycin in the presence of 10 μg/ml Brefeldin A for 4 hours at 37°C prior to live/dead staining and fixation. CD4^+^ T helper cell subsets were identified using the antibodies listed above and the addition of anti-IL-5 (for Th2) and anti-IFNγ (for Th1) after permeabilization (Invitrogen). Gates were set according to Fluorescence Minus One (FMO) controls. Representative gating strategies are shown in **Figure S1** and antibody information is listed in **Table S1.**

### Statistical analysis

All data are presented as mean ± standard error of the mean (SEM). Statistical analysis was performed using GraphPad Prism 8.0 (GraphPad Software, La Jolla, CA, USA) with unpaired t-test or one-way analysis of variance (ANOVA) followed by Fisher’s post-hoc test. Differences between groups were considered statistically significant at P < 0.05. Outliers were identified according to the two-standard deviation method.

## Results

### *Fasciola hepatica* molecules modulate Th1/Th2 skewing by LPS-stimulated dendritic cells

In order to determine the immunomodulatory effects of *F. hepatica-derived* native molecules, we first tested the impact of both somatic antigens (*fh*SA) and excretory-secretory products (*fh*ESPs) using an *in vitro* model of human monocyte-derived dendritic cells (moDCs, **Fig. 1a).** We showed that stimulation with both *fh*SA and *fh*ESPs potently induced IL-10 production in LPS-activated moDCs while neither IL-12p70 and IL-6 production nor the cell-surface expression of the main co-stimulatory markers were affected **(Fig. 1b-c,e-f).** In addition, we found that *fh*SA, and to a lesser extent *fh*ESPs, conditioned moDCs to promote Th2 skewing and concomitantly reduce Th1 polarization, ultimately leading to an increased Th2-on-Th1 ratio **(Fig. 1d,g).** We next aimed to identify the specific molecule(s) present in *fh*ESPs that drive this effect. We focused on cathepsin and Fatty Acid Binding Protein (FABP) families as they are the most abundant molecules present in adult *fh*ESPs [27]. Using a yeast expression system, we generated recombinant molecules for three of the *fh*ESP molecules, namely *fh*CL5, *fh*FABP1 and *fh*FABP5, and tested their immunomodulatory properties in moDCs **(Fig. 2a).** Although *fh*CL5 and *fh*FABP5 had only a marginal impact on moDC cytokine production and T cell skewing, *fh*FABP1 reduced production of inflammatory cytokines and potently induced stimulation of IL-10 secretion by moDCs **(Fig. 2b).** Furthermore, *fh*FABP1 instructed moDCs to promote Th2 over Th1 polarization, leading to an increase in the Th2-on-Th1 ratio **(Fig. 2c).**

**Figure 1.**
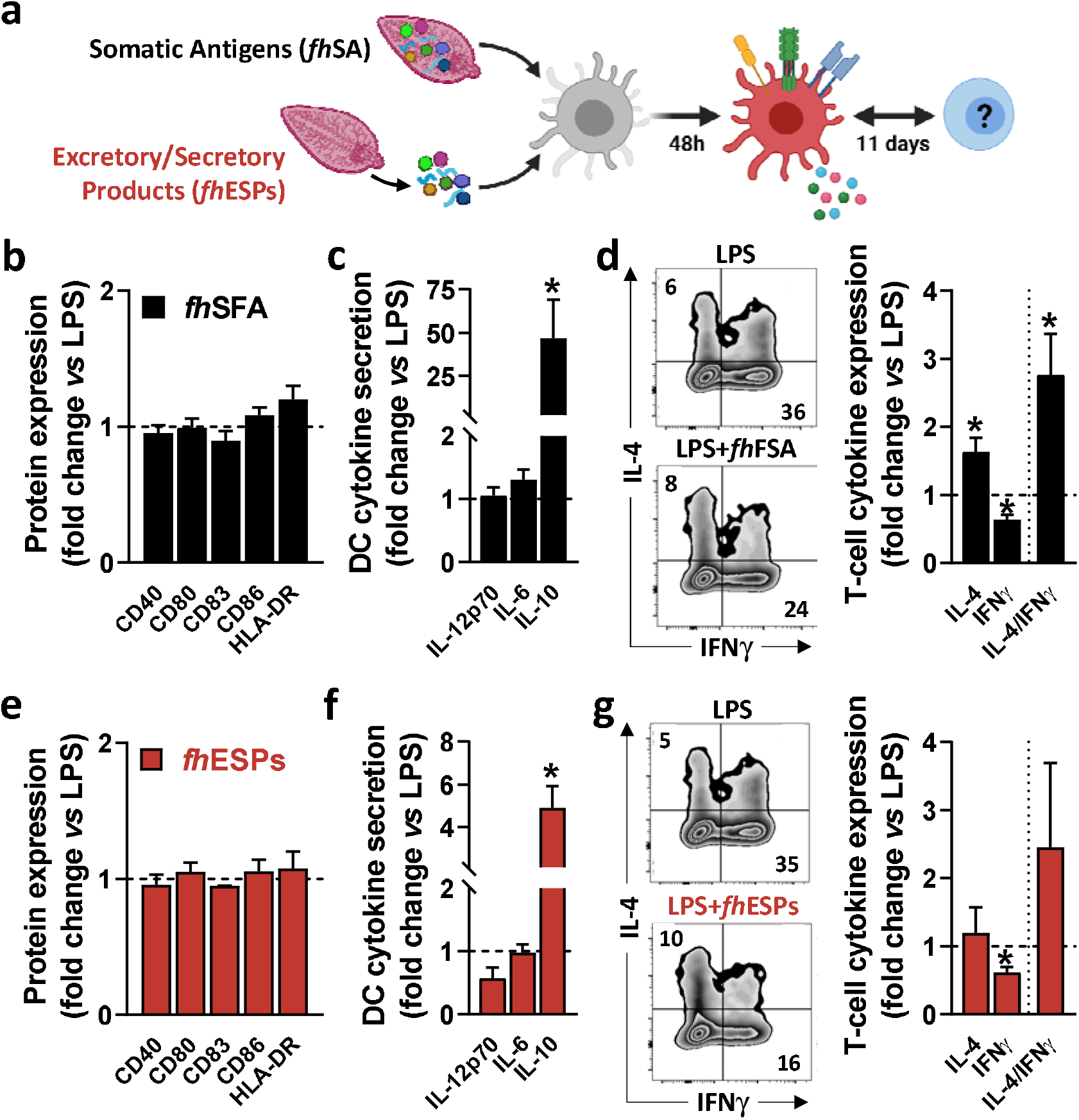
Native *Fasciola hepatica* molecules modulate Th1/Th2 skewing by LPS-stimulated dendritic cells. Human monocyte-derived DCs (moDCs) were treated with or without *F. hepatica* somatic antigens (*fh*SFA, 25 μg/ml) or excretory-secretory products (*fh*ESPs) in the presence of LPS (100 ng/ml) **(a).** After 48 hours, the cell-surface expression of co-stimulatory molecules **(b, e)** and cytokines production by moDCs **(c, f)** were determined by flow cytometry and ELISA, respectively. moDCs primed with or without *fh*SFA and *fh*ESPs were cultured with allogeneic naïve CD4^+^ T cells for 11 days in the presence of SEB and IL-2. Intracellular IL-4 and IFNγ expression was assayed by flow cytometry 6 hours after restimulation with PMA and ionomycin **(d, g),** and the IL-4-on-IFNγ ratio calculated. One representative experiment is shown for the Zebra plot. All data are expressed relative to the DCs stimulated with LPS alone (dash line) as mean ± SEM. * P ≤ 0.05 vs LPS alone (n=3 independent experiments).

**Figure 2.**
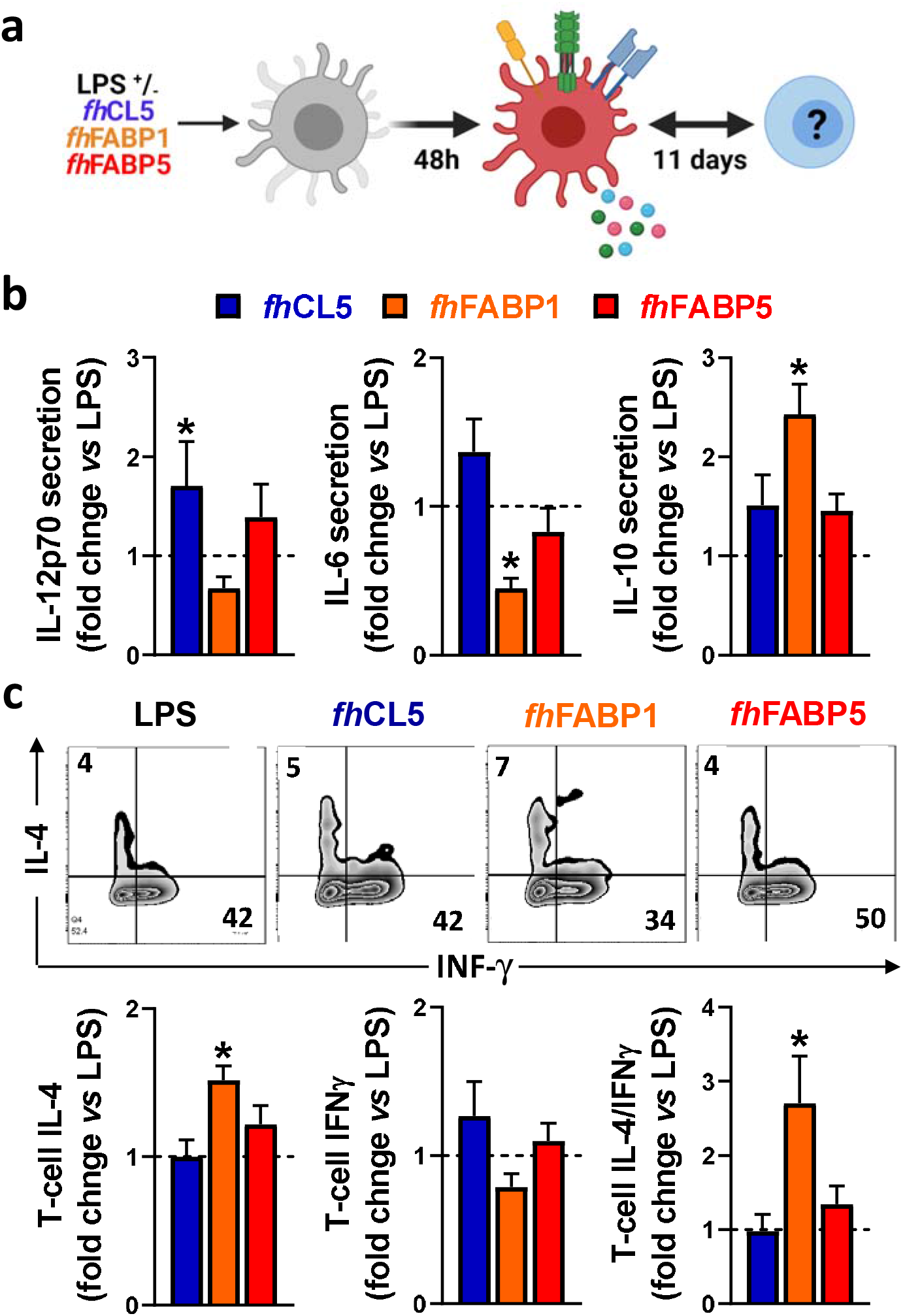
*Fasciola hepatica* FABP1, but not CL5 and FABP5, conditions dendritic cells to prime different T cell subsets. Human monocyte-derived DCs (moDCs) were treated for 48 h with or without 25 μg/ml recombinant *Fasciola hepatica* cathepsin L5 (*fh*CL5, blue bars), Fatty Acid Binding Protein 1 (*fh*FABP1, orange bars) or Fatty Acid Binding Protein 5 (*fh*FABP5, red bars) in the presence of LPS (100 ng/ml) **(a)**. Conditioned moDCs were characterized by cytokine production after 48 hours **(b)** and their capacity to prime CD4^+^ T cell responses **(c)** after 11 days of co-culture, as described in Figure 1. All data are expressed relative to the DCs stimulated with LPS alone (dash line) as mean ± SEM. * P ≤ 0.05 vs LPS alone (n=3 independent experiments).

### *Fasciola hepatica* FABP1 induces a tolerogenic-like phenotype in dendritic cells and increases both the Th2-on-Th1 ratio and IL-10-producing T cells

We next explored in more depth the impact of *fh*FABP1 on human moDCs, first showing that the molecule can bind **(Fig. 3a)** and then be rapidly internalized by the cell **(Fig. 3b** and **Fig. S2).** LPS-stimulated moDCs were next treated with increasing concentrations of *fh*FABP1 followed by determination of cytokine secretion, following either direct stimulation or secondary restimulation with CD40L, and of T cell skewing capacity **(Fig. 3c).** We confirmed that *fh*FABP1 neither induced apoptosis/cell death **(Fig. S3),** nor affected the expression of canonical DC co-stimulatory markers **(Fig. 3d),** but significantly reduced the production of pro-inflammatory cytokines IL-12p70 and IL-6, and increased IL-10 secretion by moDCs **(Fig. 3e).** When co-cultured with a CD40L-expressing cell line to mimic the interaction with CD4^+^ T cells, we also found a potent and dose-dependent inhibition of the pro-inflammatory cytokines while IL-10 secretion was not significantly changed **(Fig. 3f).** In addition, naïve CD4^+^ T cells co-cultured with *fh*FABP1-conditioned moDCs increased IL-4 and decreased IFNγ expression relative to controls, indicating an upward shift in the Th2-on-Th1 ratio **(Fig. 3g).** Furthermore, a significant increase in T cell IL-10 secretion was observed after re-stimulation with anti-CD3 and anti-CD28, suggesting that *fh*FABP1 may also enhance DC-mediated priming of regulatory T cells (Tregs) **(Fig. 3h).** However, we did not find changes in the T cell expression of the canonical Treg transcription factor FOXP3, while a significant increase in the Th2 transcription factor GATA3 was observed **(Fig. 3i).** Altogether, our data suggest that *fh*FABP1 may induce tolerogenic DCs (TolDCs), which are known to differ from immunogenic DCs through their expression of costimulatory molecules, pro-inflammatory cytokine secretion, and ability to prime Tregs [36]. We therefore compared the effects of *fh*FABP1 to those of Vitamin D3 (VitD3) - a well-known inducer of TolDCs [37; 38; 39] - on DC expression of several co-stimulatory and tolerogenic markers. As expected, VitD3-conditioned moDCs exhibited reduced expression of co-stimulatory markers CD80 and CD83, and a concomitant increase in CD163 and ILT3, both tolerogenic markers, while PD-L1 and CD103 remained unchanged **(Fig 4a-b).** By contrast, *fh*FABP1-conditioned moDCs did not exhibit increased expression of CD163 and ILT3 but displayed significant upregulation of CD103, a surface costimulatory molecule shown to be involved in induction of DC tolerance [40] **(Fig 4a-b).** Furthermore, IL-10 secretion was significantly increased by *fh*FABP1 compared to VitD3-conditioned moDCs, whereas TGFβ production was not affected by either stimulus **(Fig 4c-d).** Of note, similar features were observed when moDCs were stimulated with TNF and IL1β **(Fig. S4),** indicating that the immunomodulatory effects of *fh*FABP1 are not only mediated by LPS-induced TLR4 signalling.

**Figure 3.**
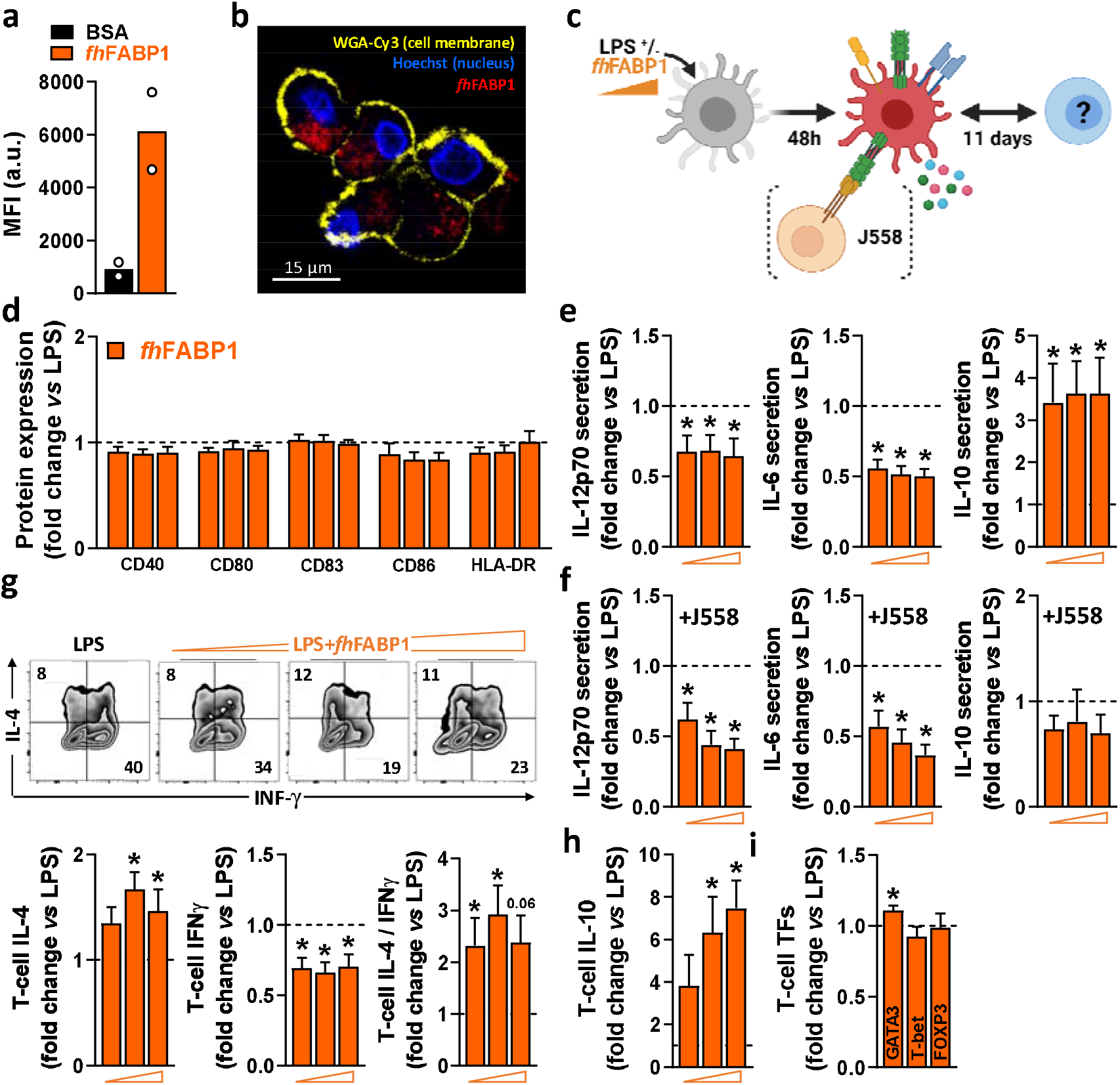
*Fasciola hepatica* FABP1 reduces the inflammatory phenotype of LPS-stimulated dendritic cells and increases both Th2-on-Th1 ratio and IL-10-producing T cells. The binding of PF-647-labeled BSA-control (black bar) or recombinant *fh*FABP1 (orange bars) to monocyte-derived DCs (moDCs) was assessed after 2 hours by flow cytometry **(a).** The subcellular localization of PF-647-labeled recombinant *fh*FABP1 (depicted in red) in moDCs was determined after 4 hours by confocal microscopy and shown as 2D images of Z-planes with nuclear (Hoechst, depicted in blue) and membrane (WGA-Cy3, depicted in yellow) staining **(b)**. One representative experiment is shown from 3 independent experiments. Human moDCs were treated for 48 h with increasing concentration of *fh*FABP1 (10, 25, 50 μg/ml) in presence of LPS (100 ng/ml) and either co-cultured with CD40-L-expressing cell line J558 for 24 hours or co-cultured with naïve CD4^+^ T cells for 11 days **(c).** The cell-surface expression of co-stimulatory molecules **(d)** and cytokines secretion by moDCs with **(e)** or without co-culture with J558 **(f)** were determined by flow cytometry and ELISA, respectively. Conditioned moDCs were characterized for their capacity to prime CD4^+^ T cell responses, as described in Figure 1 **(g).** One representative experiment is shown for the Zebra plot. In addition, primed CD4^+^ T cells were restimulated with α-CD3 and α-CD28 to measure IL-10 production after 24 hours **(h)** and analysed for GATA3, T-bet, and FOXP3 transcription factor expression by flow cytometry **(i).** All data are expressed relative to the DCs stimulated with LPS alone (dash line) as mean ± SEM. * P ≤ 0.05 vs LPS alone (n=3-7 independent experiments).

**Figure 4.**
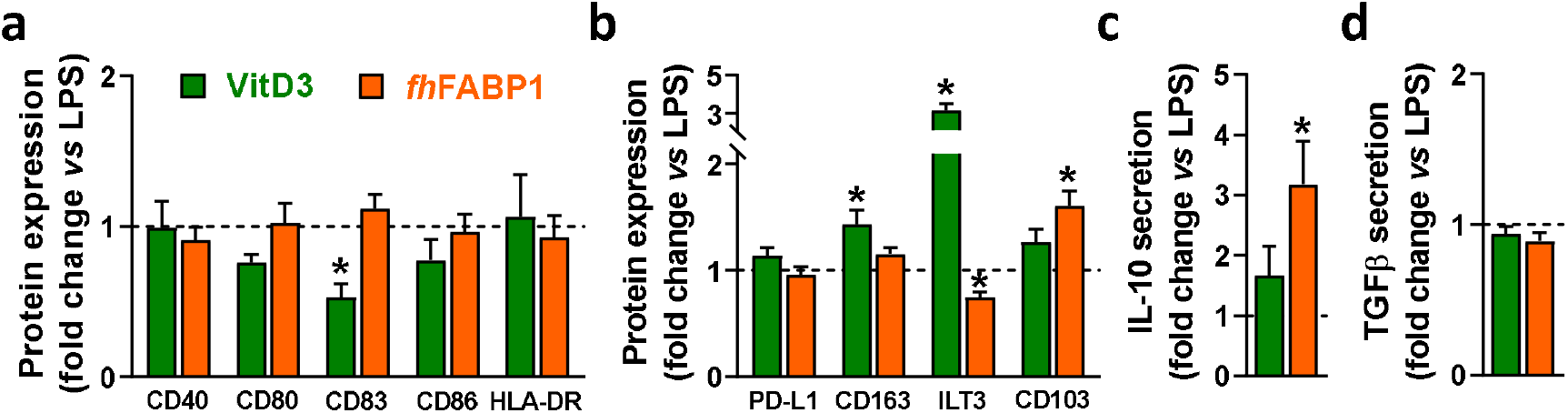
*Fasciola hepatica* FABP1 induces CD103 tolerogenic marker in dendritic cells. Human monocyte-derived DCs (moDCs) were treated with recombinant *fh*FABP1 (25 μg/ml, orange bars) or Vitamin D (VitD3, 10 nM, green bars) in the presence of LPS (100 ng/ml), as described in Figure 1. After 48 hours, the cell-surface expression of co-stimulatory molecules **(a)** and tolerogenic markers **(b)** were assessed by flow cytometry, and IL-10 and TGFβ secretion by moDCs **(c-d)** were determined by ELISA. All data are expressed relative to the DCs stimulated with LPS alone (dash line) as mean ± SEM. * P ≤ 0.05 vs LPS alone (n=4 independent experiments).

### *Fasciola hepatica* FABP1 promotes thrombospondin 1 expression and secretion by dendritic cells

In order to elucidate the underlying molecular mechanism(s) by which *fh*FABP1 modulates DC-mediated T cell polarization, we performed an in-depth DC phenotyping through unbiased analyses of cell transcriptome, lipidome and secretome, as well as intrinsic core metabolic pathways. Using RNA sequencing, we found that *fh*FABP1 induced a significant up- and downregulation of 167 and 68 transcripts, respectively, in LPS-stimulated moDCs **(Fig. 5a).** Gene ontology analysis indicated an over-representation of upregulated genes involved in immune response, cell proliferation and leukocyte chemotaxis, and of downregulated genes belonging to various type of inflammatory response pathways **(Fig. 5b).** Among the genes that were both highly expressed and the most significantly regulated by *fh*FABP1, we found some encoding secreted factors known to be involved in DC-mediated Th1 and Treg priming [41; 42; 43; 44; 45], notably *IL10*, *CXCL9*, *CXCL11*, *CCL19*, and *THBS1* (Thrombospondin 1, TSP-1) **(Fig. 5c** and **Fig. S5a).** Using a cytokine array covering more than 100 secreted proteins, we confirmed that *fh*FABP1 potently induces TSP-1 secretion by moDCs and a concomitant decrease in CXCL9, CXCL11 and CCL19 production **(Fig. 5d).** The effects of *fh*FABP1 on the two most up-(TSP-1) and downregulated (CXCL11) secreted proteins by moDCs were also confirmed by ELISA **(Fig. 5e** and **Fig. S5b).** Since changes in cellular metabolism underpins DC immune functions [36], we also assessed the impact of *fh*FABP1 on DC intrinsic core metabolic pathways using different approaches. At the transcriptional level, gene set enrichment analysis did not reveal any significant overrepresentation of glucose, fatty acid and amino acid metabolic pathways in *fh*FABP1-conditioned moDCs **(Fig. S6a).** In line with this, mitochondrial mass assessed by Mitotracker Green staining **(Fig. S6c)** and both mitochondrial oxygen consumption and glycolytic rates measured by Seahorse flux analyser **(Fig. S6b)** were not changed by *fh*FABP1 in LPS-stimulated moDCs. Furthermore, neither neutral lipids **(Fig. S6d)** or intracellular levels of a broad range of lipid species **(Fig. S6d)** were affected, indicating that *fh*FABP1 immunomodulatory effects are unlikely to be explained by modulation of DC metabolism.

**Figure 5.**
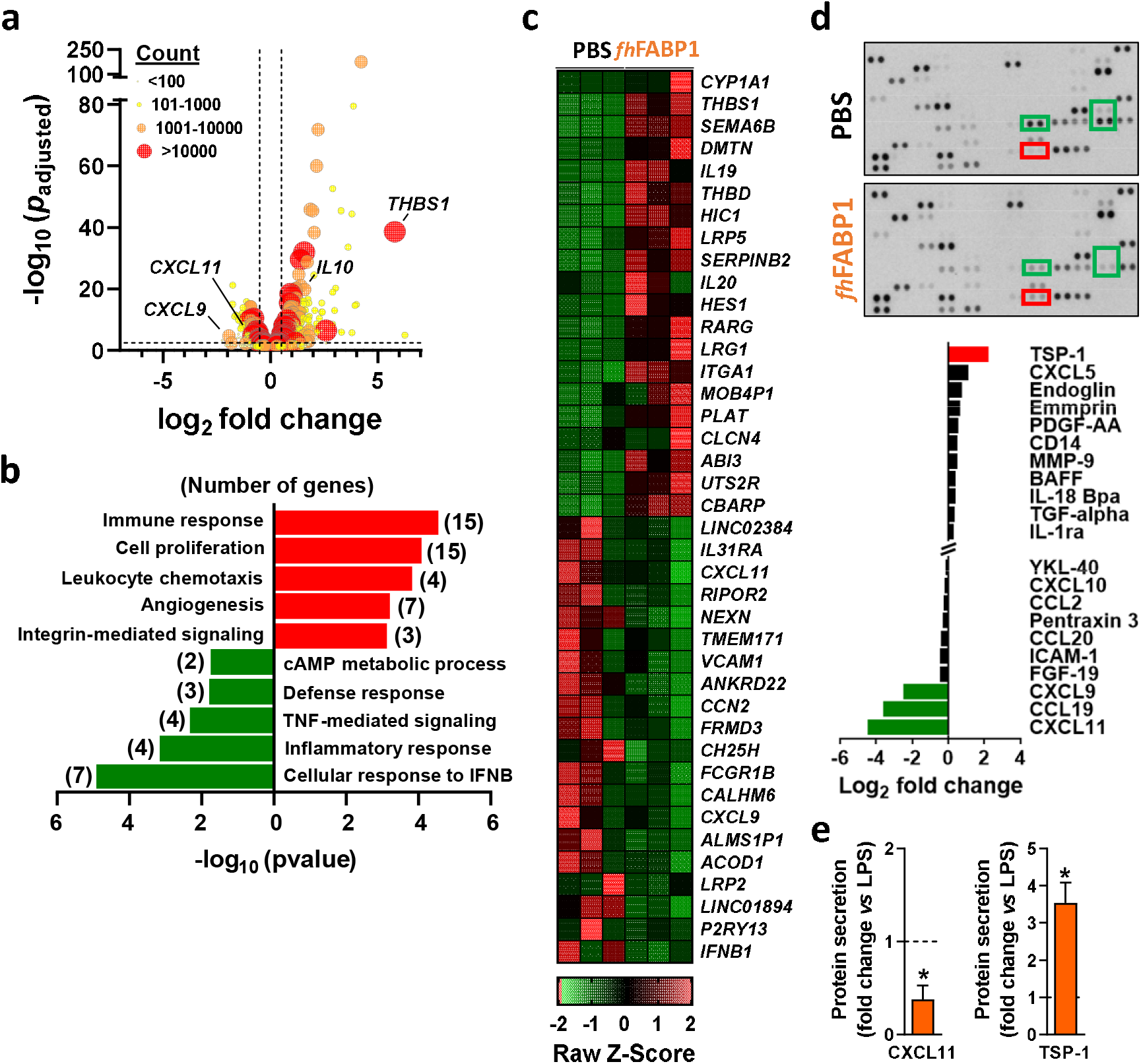
*Fasciola hepatica* FABP1 promotes thrombospondin expression and secretion by dendritic cells. Human monocyte-derived DCs (moDCs) were treated with or without recombinant *fh*FABP1 (25 μg/ml) for 5 h **(a-d)** or 48 hours **(e-f)** in the presence of LPS (100 ng/ml). RNAseq was performed in 5h-conditioned moDCs (n=3 independent experiments) and the differentially expressed genes (DEGs) are visualized on a volcano plot using DESeq2 normalized transcript counts **(a).** Gene ontology analysis was performed on the up and down DEGs **(b).** The top 20 up and down DEGs are shown on the heat map **(c).** Proteomic analysis of the moDCs secretome was performed in LPS-stimulated moDCs treated with or without recombinant *fh*FABP1 for 48 hours using the Proteome Profiler Cytokine Kit **(d).** The effects of *fh*FABP1 on secretion of CXCL11 and thrombospondin (TSP-1) by moDCs were confirmed by ELISA **(e)** and the data are expressed relative to the moDCs stimulated with LPS alone (dash line) as mean ± SEM. * P ≤ 0.05 vs LPS alone (n=3-5 independent experiments).

### *Fasciola hepatica* FABP1 modulates dendritic cell-mediated T cell polarization in a thrombospondin 1-dependent manner

We next investigated the role played by the secreted factors identified using our transcriptomic and proteomic approaches in T cell polarization by *fh*FABP1-conditioned moDCs. For this purpose, we either replenished the chemokines found to be reduced by *fh*FABP1 (CXCL11, CCL9 and CXCL9) or neutralized DC-secreted TSP-1 using a blocking antibody during DC-T cell co-culture **(Fig. 6a-b).** Although the chemokines mixture did not affect the capacity of *fh*FABP1-conditioned moDCs to modulate the Th2/Th1 balance and T cell IL-10 production **(Fig. 6c),** neutralizing TSP-1 abolished the effects of *fh*FABP1 on DC-mediated T cell polarization **(Fig. 6d).** Altogether, we demonstrate that *fh*FABP1 modulates T cell polarization by promoting TSP-1 secretion in moDCs. Finally, we assessed whether *fh*FABP1-conditioned moDCs could prime regulatory T cells with suppressive capacity, by measuring their respective ability to suppress proliferation of bystander T cells. We found that in contrast to VitD3, T cells primed by *fh*FABP1-conditioned moDCs did not inhibit the proliferation of bystander T cells **(Fig. 6e),** indicating that despite inducing IL-10-secreting T cells, *fh*FABP1 did not promote DC-mediated skewing of classical Tregs with suppressive capacity.

**Figure 6.**
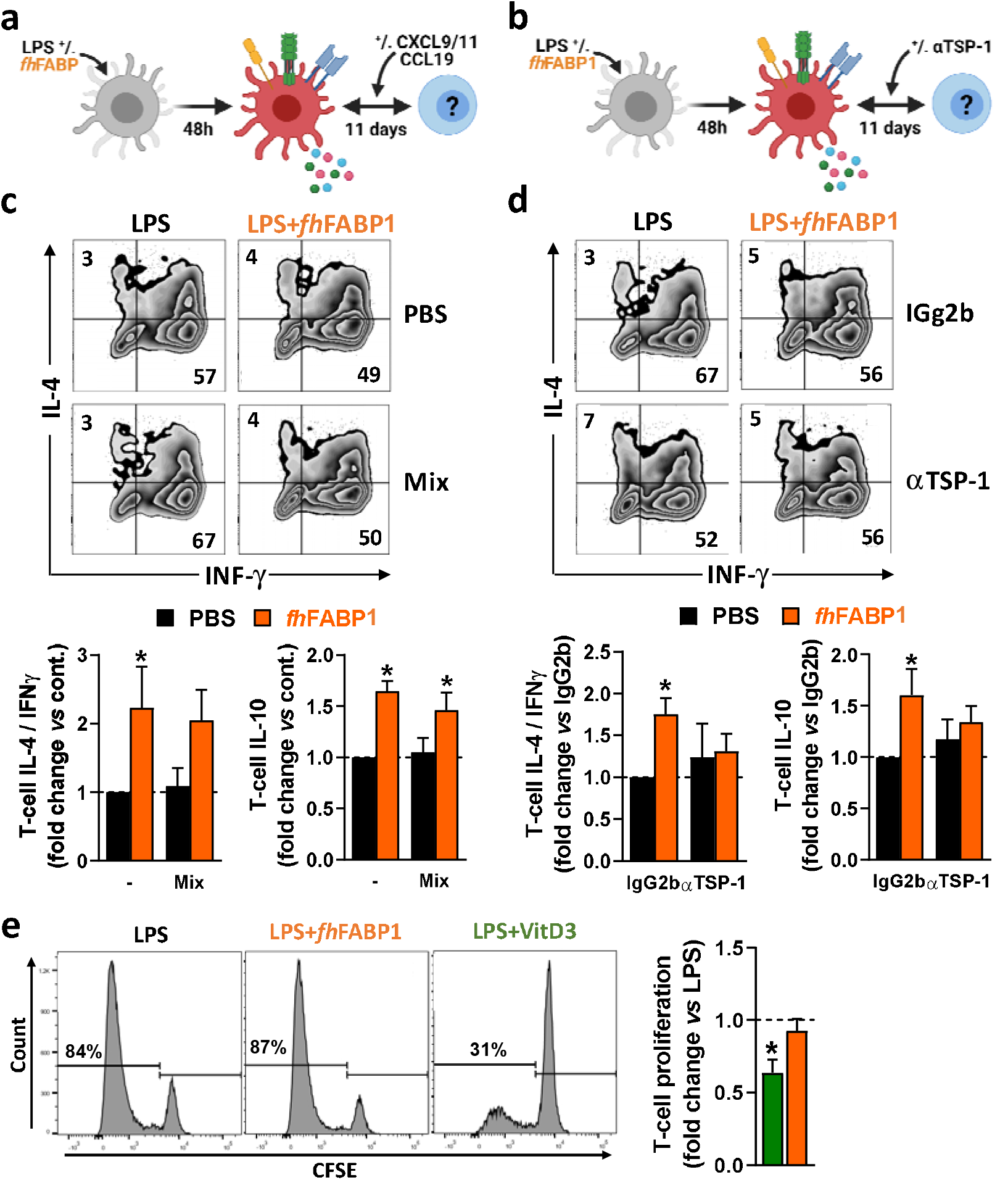
*Fasciola hepatica* FABP1 modulates dendritic cell-mediated T cell polarization in a thrombospondin 1-dependent manner but does not induce classical regulatory T cells. Human monocyte-derived DCs (moDCs) were treated with or without recombinant *fh*FABP1 (25 μg/ml) for 48 hours and then co-cultured for 11 days with allogeneic naïve CD4^+^ T cells in the presence of a chemokine mix (CXCL9, CCL9 and CXCL11 at 0.33 ng/ml each) **(a)** or αTSP-1 blocking antibody and IgG2b isotype control (10 μg/ml) **(b).** Intracellular IL-4, IFNγ and IL-10 expression was assayed by flow cytometry 6 h after restimulation with PMA and ionomycin, and the IL-4-on-IFNγ ratio calculated **(c-d).** One representative experiment is shown for the Zebra plot. The suppressive capacity of T cells induced by *fh*FABP1- or VitD3-conditioned DCs as described in Figure 4 was assessed by a T cell suppression test **(e).** One representative experiment from 4 independent experiments is shown for the histogram plots. All data are expressed relative to the DCs stimulated with LPS alone (dash line) as mean ± SEM. * P ≤ 0.05 vs LPS alone (n=3-4 independent experiments).

### *Fasciola hepatica* FABP1 induces a Th2 immune response *in vivo*, but does not affect metabolic homeostasis in obese mice

To assess whether *fh*FABP1 can also modulate T cell polarization *in vivo*, we administered the recombinant molecule, with or without LPS, via hock injection in the mouse, and then analysed T cells from the draining popliteal lymph nodes (pLNs) after 6 days **(Fig. 7a).** As expected, LPS induced a significant increase in both total pLN immune cells and CD44^+^CD4^+^ T cells, while the percentages of total CD4^+^ T cells in pLN cells remained unchanged regardless of the conditions **(Fig. 7c-d).** Remarkably, *fh*FABP1 significantly increased the percentage of IL-4-expressing CD4^+^ T cells as well as IL-4 secretion by pLN cells after *ex vivo* restimulation, while a trend for decreased IFNγ-expressing CD4^+^ T cells and IFNγ secretion by pLN cells was observed **(Fig. 7e-f).** Altogether, and in line with our *in vitro* data, *fh*FABP1 can also affect the Th2/Th1 balance towards an enhanced Th2 immune response *in vivo* **(Fig. 7e-f).**

**Figure 7.**
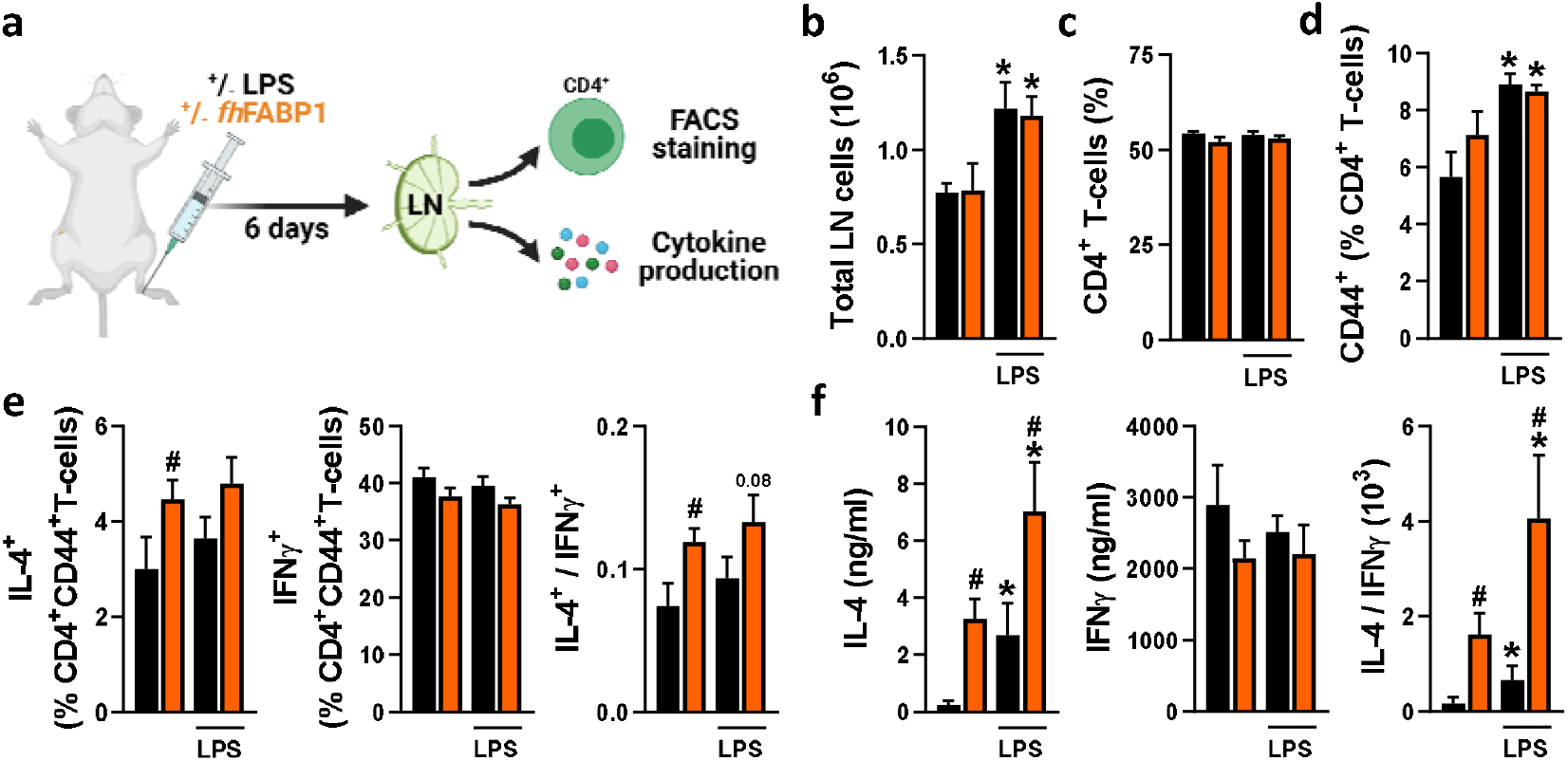
*Fasciola hepatica* FABP1 modulates T cell polarization *in vivo*. Mice were injected s.c. into the hind hock with PBS (black bars) or 25 μg recombinant *fh*FABP1 (orange bars) together with or without LPS (200 ng). The draining popliteal lymph nodes (LNs) were collected after 6 days for FACS analyses and *ex vivo* restimulation of LN cells **(a).** The total LN cells count **(b)** and the frequencies of CD4^+^ T cells **(c)** and CD4^+^CD44^high^ effector T cells **(d)** were determined. The percentages of IL-4^+^, INFγ^+^ and IL-10^+^ cells within the CD4+CD44^high^ effector T cell population were also assessed, and the IL-4-on-IFNγ ratio plotted **(e).** Alternatively, part of the LN cells were re-stimulated *ex vivo* and secretion of IL-4 and IFNγ were measured by cytokine bead assay **(e).** Results are expressed as mean ± SEM. * P ≤ 0.05 vs LPS alone, # P ≤ 0.05 vs PBS (n=3-5 mice per group).

Infection with various helminth species or treatment with their immunomodulatory molecules have been reported to restore type 2 immunity in metabolic organs and improve insulin sensitivity and glucose homeostasis in obese mice [6]. Therefore, we investigated whether *fh*FABP1-induced Th2 immune response may alleviate obesity-induced metabolic dysfunctions. For this purpose, HFD-fed mice received biweekly intraperitoneal injections with *fh*FABP1 for four weeks **(Fig. 8a),** a treatment that did not affect body weight gain and body composition in obese mice **(Fig. 8b-d).** Immune cell phenotyping in metabolic tissues showed that although no differences in Kupffer cell and CD4^+^ T cells numbers were observed in the liver **(Fig. S7),** *fh*FABP1 markedly increased both total leukocytes and CD4^+^ T cells in white adipose tissue (WAT) **(Fig. 8e-f).** Strikingly, *fh*FABP1 potently increased the numbers of CD44^+^ CD4^+^ T cells (*data not shown*), which were characterized by increased frequencies of IL-5^+^ Th2 cells, and by a trend for decreased IFNγ^+^ Th1 cells **(Fig. 8g-h).** This was in line with what was observed in draining pLNs after hock injection of *fh*FABP1. Furthermore, the numbers of eosinophils **(Fig. 8i)** and YM1^+^ macrophages were also increased while total macrophages and RELMα^+^ macrophages were unchanged **(Fig. 8j-k).** However, this potent WAT type 2 immune response was not associated with any beneficial effects on HFD-induced metabolic dysfunctions, since neither fasting glycemia, nor whole-body insulin sensitivity and glucose tolerance were improved by *fh*FABP1 treatment **(Fig. 8l-m).**

**Figure 8.**
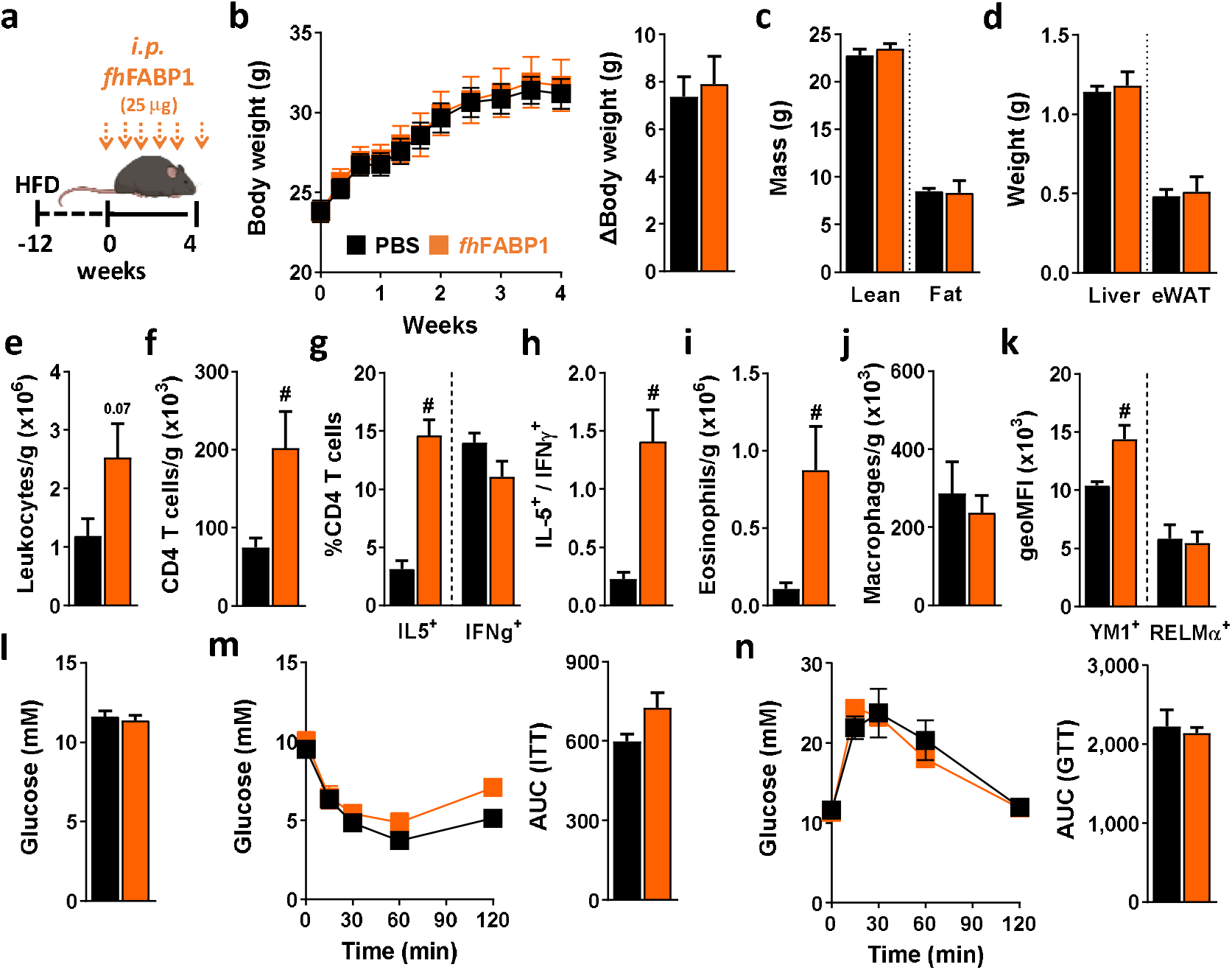
*Fasciola hepatica* FABP1 induces a type 2 immune response in adipose tissue but without affecting metabolic homeostasis in obese mice. Mice were fed a high-fat diet (HFD) for a period of 12 weeks and then received intraperitoneal injections of 25 μg recombinant *fh*FABP1 (orange squares/bars) or vehicle (PBS, black squares/bars) every 3 days for 4 weeks **(a).** Body weight was monitored throughout the supplementation period **(b).** At week 4, body composition was determined **(c).** After sacrifice, the weights of the liver and eWAT were measured **(d).** The number of eWAT total leukocytes **(e),** CD4^+^ T cells **(f),** eosinophils **(i)** and total macrophages **(j),** together with the percentage of CD44^+^, IL-5^+^, IFNγ^+^ CD4^+^ T cells **(g)** and the YM1 and RELMα mean expression on macrophages **(k)** were determined by flow cytometry. The IL-5^+^-on-IFNγ^+^ ratio was calculated **(h).** At week 4, fasting blood glucose levels were measured **(l).** At week 3-4, whole-body insulin sensitivity **(m)** and glucose tolerance **(n)** were performed and the respective AUC was calculated. Results are expressed as mean ± SEM. # P ≤ 0.05 *vs* HFD (n=5-6 mice per group).

Altogether, we showed that although *fh*FABP1 modulates T cell polarization, notably by promoting DC TSP-1 secretion *in vitro*, it does not affect metabolic homeostasis in a mouse model of obesity and type 2 diabetes.

## Discussion

The parasitic trematode *F. hepatica* evades the host immune defense mechanisms through secretion of various specific immunomodulatory molecules that help to manipulate the immune response, ultimately allowing maintenance of long-term infection. FABPs are among the parasite main excreted/secreted proteins and have been shown to display some antiinflammatory properties, notably in macrophages [30]. However, little was known regarding FABPs impact on DCs and their subsequent capacity to prime specific CD4^+^ T cell subsets. In the present study we showed that *fh*FABP1 modulates T cell polarization *in vitro* by promoting thrombospondin-1 secretion by DCs, an effect which is also observed *in vivo* but that is not associated with improvements of whole-body metabolic homeostasis in obese mice.

The characterization of the T cells primed by *fh*FABP1-conditioned human moDCs in our *in vitro* co-culture model revealed an increase in both Th2-on-Th1 ratio and IL-10-secreting T cells. Although a significant rise in T cell expression of the Th2-specific transcription factor GATA3 was evidenced, there was no change in the expression of the canonical regulatory T cell transcription factor FOXP3, suggesting that *fh*FABP1-conditioned DCs may induce FOXP3^-^IL-10^+^ Tr1 cells. Interestingly, studies investigating T cell responses following various helminth infections have previously reported induction of both classical FOXP3^+^IL-10^+^ and non-conventional FOXP3^-^IL-10^+^ regulatory T cells [46]. Furthermore, *F. hepatica-specific* Tr1-cell lines generated from infected mice have also been shown to suppress T cell proliferation and IFNγ production by Th1 cells [25]. Despite Tr1 cells are usually known to exert regulatory activity [47], we did not find any inhibition of effector T cells proliferation by *fh*FABP1-induced T cells. Although we cannot exclude species-specific properties, the exact identity and function of these *fh*FABP1-induced FOXP3^-^IL-10-producing T cells remains to be elucidated. Of note, among the limited data available, human moDCs stimulated with *F. hepatica* somatic antigens were reported to inhibit proliferation of CD4^+^ allogeneic T cells, although both the underlying mechanism and functional characterization of the T cells were not investigated [26].

When investigating more in depth the mechanism(s) involved in the polarization of naïve T cells by *fh*FABP1-conditioned human moDCs, we found no impact on DC expression of co-stimulatory molecules and HLA-DR, while significantly lower levels of pro-inflammatory cytokines IL12p70 and IL-6 and an increased level of IL-10 were produced. Despite no obvious effect on maturation, these cells displayed some features of tolerogenic DCs [48; 49; 50; 51]. Nevertheless, the analysis of cell-surface expression of canonical tolerogenic markers only revealed a significant increase in CD103. Of note, CD103 expression on human DCs has been shown to be triggered by retinoic acid [52], which has been reported itself as a potential ligand for native *F. hepatica* FABPs [53]. It is well acknowledged that modulation of intrinsic cellular metabolism shapes the functional properties of DCs to drive polarization of different Th cell subsets by orchestrating the engagement of distinct metabolic pathways [54]. As such, tolerogenic-like DCs have been suggested to be mainly dependent on fatty acid oxidation and mitochondrial oxidative metabolism [36; 55]. However, in our experimental conditions, we found no obvious impact of *fh*FABP1 on both cellular glycolysis and mitochondrial oxidative phosphorylation, indicating that DC-mediated priming of Th2 and IL-10-producing T cells by *fh*FABP1 occurs independently of metabolic perturbations. To get some insights on the putative mechanism by which *fh*FABP1 modulate DC-mediated T cell polarization, we performed an unbiased analysis of the DC transcriptome and measured a large portfolio of their secreted factors. Remarkably, we identified TSP-1 as a key player involved in DC-mediated priming of Th2 and IL-10 producing T cells by *fh*FABP1. TSP-1 is indeed a potent anti-inflammatory chemokine and its interaction with the T cell receptor CD47 has been shown to be involved in Treg differentiation [45; 56]. TSP-1 signaling through CD47 inhibits TCR signaling via several mechanisms, including modulation of intracellular calcium homeostasis, cyclic nucleotide signaling and nitric oxide biosynthesis [57]. In addition, TSP-1 has also been reported to act as an autocrine negative regulator of DC activation that might contribute to DC exhaustion [44]. While this aspect of the DC-T cell regulation requires further extensive investigation, we showed that *fh*FABP1 increases Th2-on-Th1 ratio and FOXP3^-^L-10-producing T cells by promoting TSP-1 secretion. Interestingly, during the preparation of our manuscript, a study reported the immunomodulatory effects of native *fh*FABP1 (named Fh12 by the authors) on mouse bone marrow-derived DCs (BMDCs) [58]. In line with our findings, this study also showed that *fh*FABP1 reduced secretion of IL-6 and IL12p70 and increased IL-10 secretion by LPS-stimulated BMDCs, resulting in a subsequent decrease in Th1 and promotion of Th2/Treg development [58]. However, DC-mediated TSP-1 production was not evaluated and the authors attributed most of the *fh*FABP1 immunomodulatory effects to induction of apoptosis, a feature that is clearly not observed in our study. Although some species-and/or experimental setting-specific effects cannot be excluded, it is tempting to suggest that some contaminant molecules isolated together with native *fh*FABP1, including other FABP isoforms, may drive most of this effect. Further studies are required to clarify this discrepancy between the different findings.

It is well established that the immune system plays a central role in the regulation of metabolic homeostasis, notably by contributing to maintenance of insulin sensitivity in metabolic organs [59; 60]. Although the exact molecular mechanisms remain unclear, adipose tissue Th2 cells, type 2 innate lymphoid cells and eosinophils have been shown to produce the canonical type 2 cytokines IL-4, IL-5 and IL-13, promoting alternative activation of macrophages and tissue-specific insulin sensitivity in homeostatic healthy conditions [61; 62]. Congruent with this, an immune-dependent component is also involved in the control of hepatic insulin sensitivity [63]. These finely-tuned immunometabolic processes are impaired during obesity, where a Th1 immune environment is believed to contribute to local low-grade inflammation and metabolic dysfunctions [64; 65]. Since we showed that *fh*FABP1 induced changes in the Th2/Th1 balance, both *in vitro* and *in vivo*, we investigated whether repeated administration of the recombinant molecule could improve metabolic homeostasis in obesity-induced insulin-resistant mice. Although treatment with *fh*FABP1 clearly induced a potent type 2 immune response in eWAT, characterized by increased Th2 cells, eosinophils and YM1-expressing alternatively activated macrophages (AAMs), we did not observe any protective effect against HFD-induced metabolic dysfunctions in obese mice. Altogether, these results may challenge the current paradigm, where promoting AAMs is associated with enhanced tissue insulin sensitivity, as previously reported in response to helminth infection or helminth-derived molecules [66]. It is however important to mention that the exact mechanism(s) by which AAMs contribute to WAT insulin sensitivity is incompletely understood. One could speculate that the Th2 cells and/or AAMs induced by *fh*FABP1 in WAT are phenotypically and functionally different than the ones induced by other type of helminth molecules. Indeed, the recent advances in high dimensional analyses of *in situ* immune cells have revealed the large diversity of cell subsets, notably in adipose tissue where macrophages display a spectrum of phenotypes that are dependent on both the tissue microenvironment and local antigen-mediated stimuli [67; 68; 69]. Supporting this hypothesis, we found that while *fh*FABP1 promoted YM1 expression in WAT macrophages, the expression of another canonical AAM marker RELMα remained unchanged. Further studies are needed to resolve this discrepancy with the current paradigm, by characterizing in more depth the immune cell phenotypes induced by *fh*FABP1 *in vivo*, specifically their tissue-specific transcriptomic and proteomic signatures. Of note, a direct effect of *fh*FABP1 on adipocytes that may counteract the beneficial type 2 immune response in WAT cannot be excluded and would be interesting to investigate in future experiments.

In conclusion, we show that *fh*FABP1 modulates T cell polarization, notably by promoting thrombospondin-1 secretion by dendritic cells *in vitro*, but does not affect metabolic homeostasis in a mouse model of type 2 diabetes. Further studies on the potential of *fh*FABP1 as an anti-inflammatory molecule are needed, such as those involving other in *vivo* models of pro-inflammatory diseases including asthma, ulcerative colitis and/or multiple sclerosis. Altogether, our work contributes to an increasing understanding of the immunomodulatory functions of a large diversity of unique helminth-derived products, that remains of utmost interest as it may pave the way to the future development of new types of therapeutics to treat inflammatory disorders.

## Acknowledgements

This work was supported by funding from the Polish Ministry of Science and Higher Education project Mobilność V (DN/MOB/278/V/2017) (to AZD), the National Science Centre in Poland project PRELUDIUM 2018/29/N/NZ6/01670 (to AK), the NWO Graduate School Program 022.006.010 (to HJPvdZ), the Dutch Lung Foundation AWWA Consortium [12.0.17.001] (to HHS and TG), DON Foundation and the Dutch Diabetes Research Foundation (to AZ and BG), and the Dutch Organization for Scientific Research (ZonMW TOP Grant 91214131 (to BG) & Vidi grant 91714352 (to HHS). We thank Frank Otto and Arifa Ozir-Fazalalikhan for their precious technical help, and Dr. Bart Everts for his critical reading of the manuscript.

## Author contributions

A. Zawistowska-Deniziak: Conceptualization, Investigation, Formal analysis, Data curation, Writing - original draft, review & editing

J.M. Lambooij: Investigation, Formal analysis, Writing - review & editing

Alicja Kalinowska: Investigation, Writing - review & editing

Thiago A. Patente: Investigation, Formal analysis, Writing - review & editing

Maciej Łapiński: Investigation, Formal analysis, Writing - review & editing

Hendrik P. van der Zande: Investigation, Formal analysis, Writing - review & editing

Katarzyna Basałaj: Investigation, Writing - review & editing

Clarize de Korne: Investigation, Formal analysis, Writing - review & editing

Mathilde A.M. Chayé: Investigation, Writing - review & editing

Tom Gasan: Investigation, Writing - review & editing

Luke J. Norbury: Investigation, Writing - review & editing

Martin Giera: Investigation, Supervision, Writing - review & editing

Arnaud Zaldumbide: Supervision, Writing - review & editing

Hermelijn H. Smits: Supervision, Writing - review & editing

B. Guigas: Conceptualization, Supervision, Writing - original draft, review & editing

## Supplementary methods

### Confocal imaging

5 × 10^4^ moDC/chamber were seeded onto poly-D-lysine (Sigma-Aldrich) coated coverslips of a glass bottom dish (ø35 mm; MatTek Corporation) for 24 hours. Cells were incubated with LPS (100 ng/ml) and *fh*FABP1 (25 μg/ml) labelled with Promofluor-660 labeling kit (Promokine) for 4 hours. The nuclei of the cells were stained with Hoechst (1:250, 1 mg/ml) and the cell membrane was stained with WGA-Cy3 (1:100, 14 μM). Images were taken at room temperature on a Leica TCS (true confocal scanning) SP8 WLL (white light laser) microscope (Leica Microsystems). The sequential scanning mode was applied to image Hoechst (excitation: 405 nm; emission: 420-470 nm), WGA-Cy3 (excitation: 560 nm, emission: 570-620 nm) and PromoFluor-660-*fh*FABP1 (excitation: 647 nm; emission: 655-705 nm). For imaging the uptake of PromoFluor-660-*fh*FABP1, a 63x objective (Leica HC PL APO 63x/1.40na OIL CS2) was used. The images were generated using the Leica software (LAS X; Leica Microsystems).

### Computational analyses of RNA sequencing data

The sequencing data quality assessment was performed using FastQC v. 0.11.9 and MultiQC v. 1.7 [1]._Trimming of low-quality sequences and adapter content was performed with cutadapt v. 2.4 [2], with the following parameters: --error-rate 0.1 --quality-cutoff 20 --minimum-length 10. The alignment of reads to the GRCh38 human reference genome has been performed using STAR v. 2.7.7a [3], using Ensembl GRCh38 version 102 genome annotation. Standard ENCODE alignment options were used, namely: --outFilterType BySJout --outFilterMultimapNmax 20 --alignSJoverhangMin 8 --alignSJDBoverhangMin 1 --outFilterMismatchNmax 999 --outFilterMismatchNoverReadLmax 0.04 --alignIntronMin 20 --alignIntronMax 1000000 --alignMatesGapMax 1000000. The duplicated alignments were flagged by the MarkDuplicates program from the Picard v.2.24.0 software suite [4]. The alignment filtering was performed with the samtools v. 1.10 software [5], with the following parameters: -q 1 -f3 -F1280. The read quantification has been performed using htseq-count script from the HTSeq v. 0.13.5 python package [6]. The filtered alignments were counted per exon on gene-level, with the default “union” counting method. The data processing pipeline was written in Nexflow [7]. The source code can be accessed from the GitLab code repository: https://gitlab.com/lapinskim/pbmcs_seq. Differential expression analysis for the stimulated PBMCS cells has been performed in R programming language v. 4.0.3 [8] with DESeq2 v. 1.30.0 R package [9]. The log fold change shrinkage for visualisations was performed using the apeglm algorithm [10].

### Metabolic profiling of moDCs

For assessing their metabolic characteristics. 5 x 10^4^ moDCs stimulated with LPS (100 ng/ml) and *fh*FABP1 (25 μg/ml) for 5 hours were plated in unbuffered, glucose-free RPMI supplemented with 5% FCS and left to rest 1 hour at 37°C in an CO^2^ free incubator. Extracellular acidification rate (ECAR) and oxygen consumption rate (OCR) were next measured using a Seahorse XFe96 Extracellular Flux Analyzer (Seahorse Bioscience) before and after addition of glucose (10 mM), oligomycin (1 μM), fluoro-carbonyl cyanide phenylhydrazone (FCCP, 3 μM), and rotenone/antimycin A (1/1 μM) (all Sigma-Aldrich).

### Lipidomic analysis

moDCs (0.5 x 10^6^/ml) were stimulated with 100 ng/ml LPS and 25 μg/ml *fh*FABP1. After 48 hours cells were harvested, washed twice with PBS and snap frozen. Lipids were extracted by the methyl-tert-butylether method and analyzed using the Lipidyzer^™^, a direct infusiontandem mass spectrometry (DI-MS/MS)-based platform (Sciex, Redwood City, USA), as previously described [11]. Lipid concentrations were expressed as nmol/10^6^ cells.

## Supplementary table

**Table S1.** FACS antibodies and ELISA kits.

| Target | Type | Source | Identifier |
| --- | --- | --- | --- |
| **Human** |  |  |  |
| Aqua | Cell death marker | Invitrogen/Thermo | L34957 |
| 7-AAD | Cell death marker | Invitrogen/Thermo | A1310 |
| CD40 | Antibody | BD Pharmingen | 555591 |
| CD80 | Antibody | BD | 560442 |
| CD83 | Antibody | Invitrogen | 12-083942 |
| CD86 | Antibody | BD Pharmingen | 561128 |
| HLA-DR | Antibody | Invitrogen | 47-9956-42 |
| CD163 | Antibody | BioLegend | 333608 |
| CD103 | Antibody | BioLegend | 350211 |
| CD85k (ILT3) | Antibody | BioLegend | 333015 |
| CD274 (PD-L1) | Antibody | eBioscience | 12-5983 |
| IL-4 | Antibody | BD Bioscience | 554516 |
| IFNγ | Antibody | BD Bioscience | 560371 |
| IL-10 | Antibody | BD Bioscience | 554707 |
| FoxP3 | Antibody | eBioscience | 17-4776-42 |
| GATA3 | Antibody | eBioscience | 50-9966-42 |
| T-bet | Antibody | eBioscience | 45-5825 |
| CD3 | Antibody | eBioscience | 46-0036 |
| CD4 | Antibody | BD Bioscience | 557852 |
| CD25 | Antibody | BD Bioscience | 340907 |
| TSP-1 | Antibody | Merck/Sigma-Aldrich | MABT879 |
| IgG2b | Antibody | BioLegend | 400347 |
| CCL19 | Chemokine | PeproTech | 300-29B |
| CXCL9 | Chemokine | PeproTech | 300-26 |
| CXCL11 | Chemokine | PeproTech | 300-46 |
| IL-10 | ELISA kit | BD Bioscience | 555157 |
| IL-12p70 | ELISA kit | BD Bioscience | 555183 |
| IL-6 | ELISA kit | BD Bioscience | 555220 |
| TSP-1 | ELISA kit | R&D Systems | DY3074 |
| TGFβ | ELISA kit | R&D Systems | DY240 |
| CXCL11 | ELISA kit | R&D Systems | DY672 |
| **Mouse** |  |  |  |
| B220 | Antibody | eBioscience | 11-0452 |
| CD3 | Antibody | Biolegend | 100237 |
| CD4 | Antibody | BD | 563232 |
| CD11b | Antibody | eBioscience | 11-0112 |
| CD11c | Antibody | BD | 553801 |
| CD44 | Antibody | eBioscience | 56-0441 |
| CD44 | Antibody | eBioscience | 48-0441 |
| CD45 | Antibody | Biolegend | 103149 |
| CD64 | Antibody | Biolegend | 139304 |
| F4/80 | Antibody | Biolegend | 123147 |
| GR-1 | Antibody | BD | 553127 |
| IFNγ | Antibody | BD | 561479 |
| IFNγ | Antibody | eBioscience | 25-7311 |
| IL-4 | Antibody | eBioscience | 17-7041 |
| IL-5 | Antibody | Biolegend | 504303 |
| IL-10 | Antibody | eBioscience | 12-7101 |
| Ly6C | Antibody | Biolegend | 128025 |
| NK 1.1 | Antibody | eBioscience | 11-5941 |
| Siglec-F | Antibody | BD | 740388 |
| YM1-biotin | Antibody | R&D Systems | BAF2446 |
| Streptavidin-PerCP-Cy5.5 | Reagent | Biolegend | 405214 |
| Zombie-UV viability kit | Cell death marker | Biolegend | 423107 |

**Figure S1.**
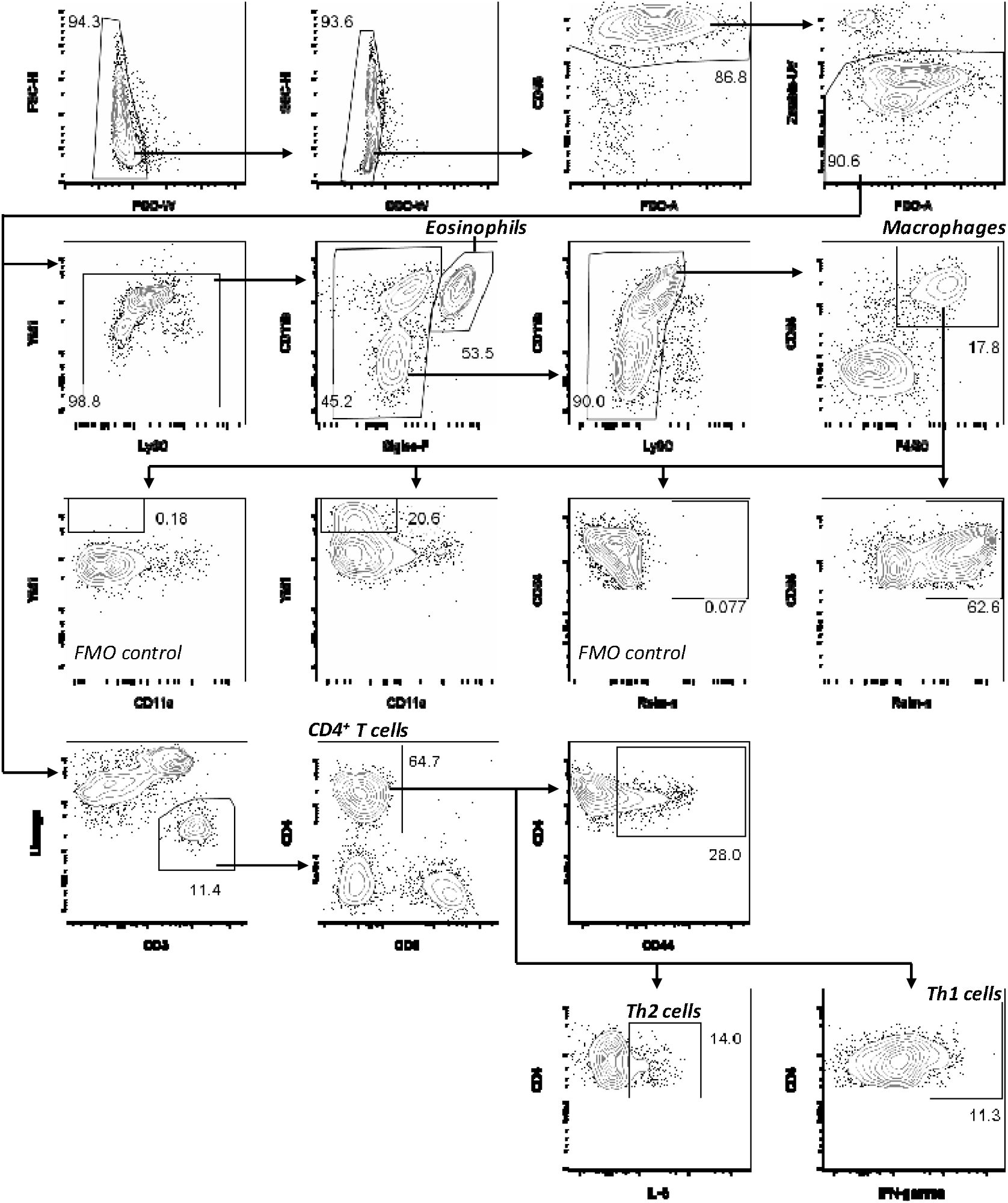
Gating strategies. Representative gating strategies for the identification of eosinophils, macrophages, Relmα^+^/CD11c^-^YM1^+^/ macrophages, CD4^+^ T cells, IL-5^+^ Th2 cells and IFNγ^+^ Th1 cells in eWAT. A sample from the *fh*FABP1-treated group is shown as example. Gating strategies were similar for liver samples.

**Figure S2.**
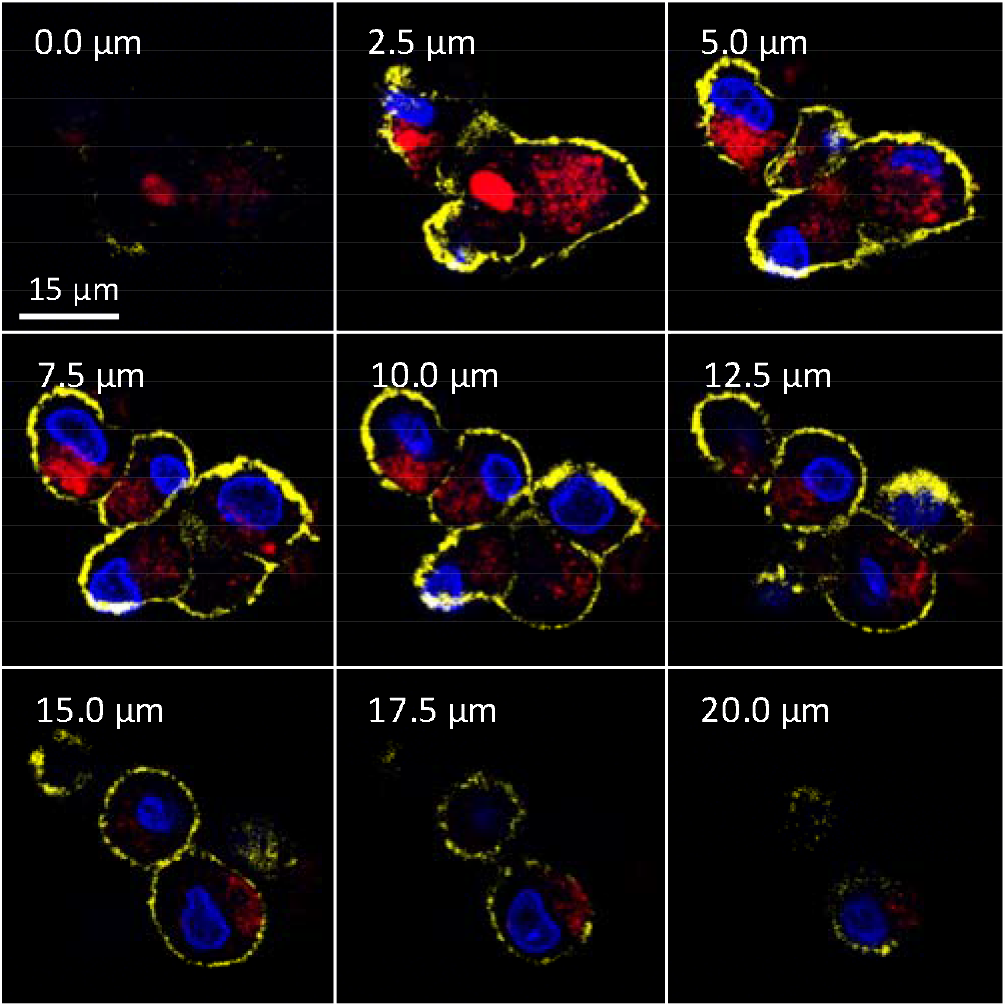
*Fasciola hepatica* FABP1 is internalized by dendritic cells and mainly localized in the intracellular compartment. The subcellular localization of PF-647-labeled recombinant *fh*FABP1 (25 μg/ml; depicted in red) in moDCs was determined after 4 hours by confocal microscopy and shown as 2D images of different Z-planes with nuclear (Hoechst, depicted in blue) and membrane (WGA-Cy3, depicted in yellow) staining. One representative experiment is shown from 3 independent experiments.

**Figure S3.**
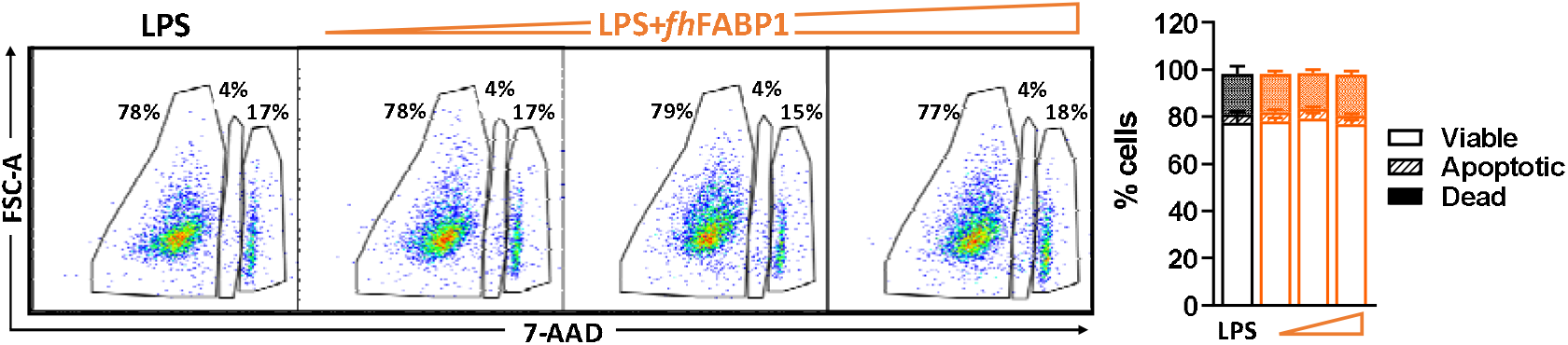
*Fasciola hepatica* FABP1 does not induce apoptosis/cell death. Human moDCs were treated for 48 h with increasing concentration of *fh*FABP1 (10, 25, 50 μg/ml) in presence of LPS (100 ng/ml). Viable, apoptotic and late apoptotic/dead cells were determined by flow cytometry as described in [12]. One representative experiment is shown for the Zebra plot. All data are expressed as mean ± SEM (n=4 independent experiments).

**Figure S4.**
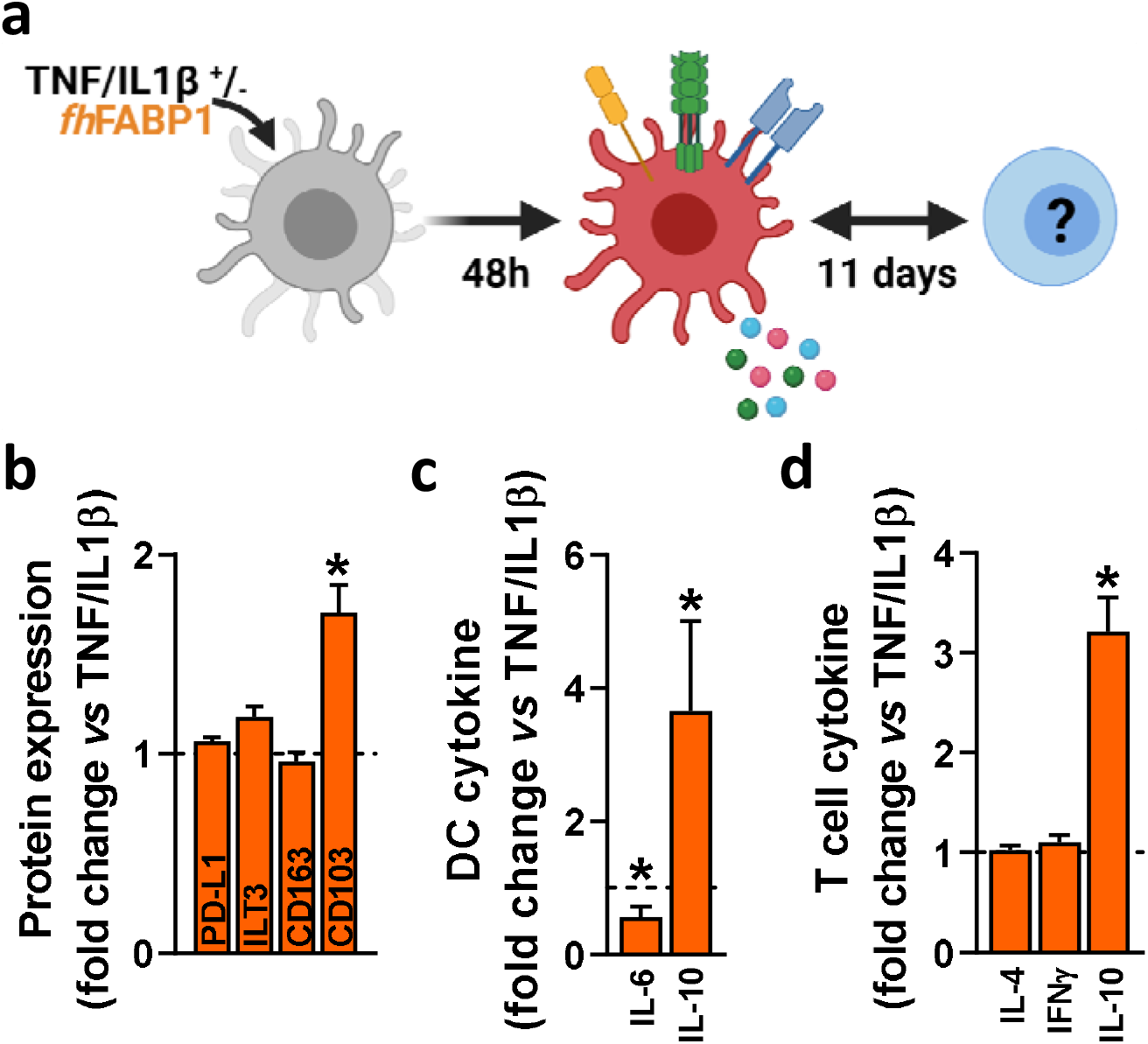
*Fasciola hepatica* FABP1 induces CD103 expression in TNFα/IL-1β-primed dendritic cells and IL-10 producing T cells. Human moDCs were treated for 48 h with or without 25 μg/ml recombinant *fh*FABP1 (orange bars) in presence of TNFα (0.05 μg/ml) and IL-1β (0.025 μg/ml) as a maturation cocktail **(a).** The cell-surface expression of tolerogenic markers **(b)** and cytokines secretion **(c)** by moDCs were determined by flow cytometry and ELISA, respectively. Conditioned moDCs were characterized for their capacity to prime CD4+ T response **(d)** after 11 days of co-culture, as described in Figure 1. All data are expressed relative to the DCs stimulated with TNFα/IL-1β alone (dash line) as mean ± SEM. * P ≤ 0.05 *vs* LPS alone (n=3-7 independent experiments).

**Figure S5.**
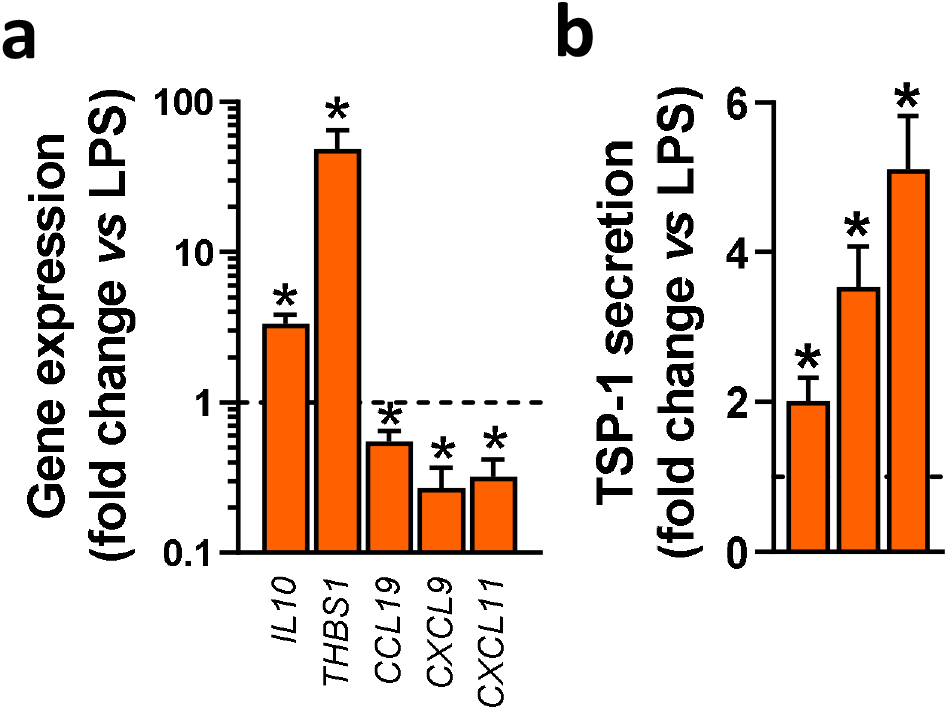
*Fasciola hepatica* FABP1 affects mRNA and protein expression of various cytokines/chemokines in LPS-stimulated dendritic cells. Human moDCs were stimulated for 5 h **(a)** or 48h **(b)** with PBS or various concentrations of recombinant *fh*FABP1 in presence of 100 ng/ml LPS. qPCR confirmation of RNAseq data showing changes in gene expression levels of various cytokines/chemokines in moDCs stimulated with 25 μg/ml *fh*FABP1 **(a).** The effect of increasing concentrations (10, 25, 50 μg/ml) of *fh*FABP1 on TSP-1 secretion by moDCs was determined by ELISA **(b).** All data are expressed relative to the DCs stimulated with LPS alone (dash line) as mean ± SEM. * P ≤ 0.05 *vs* LPS alone (n=3-5 independent experiments).

**Figure S6.**
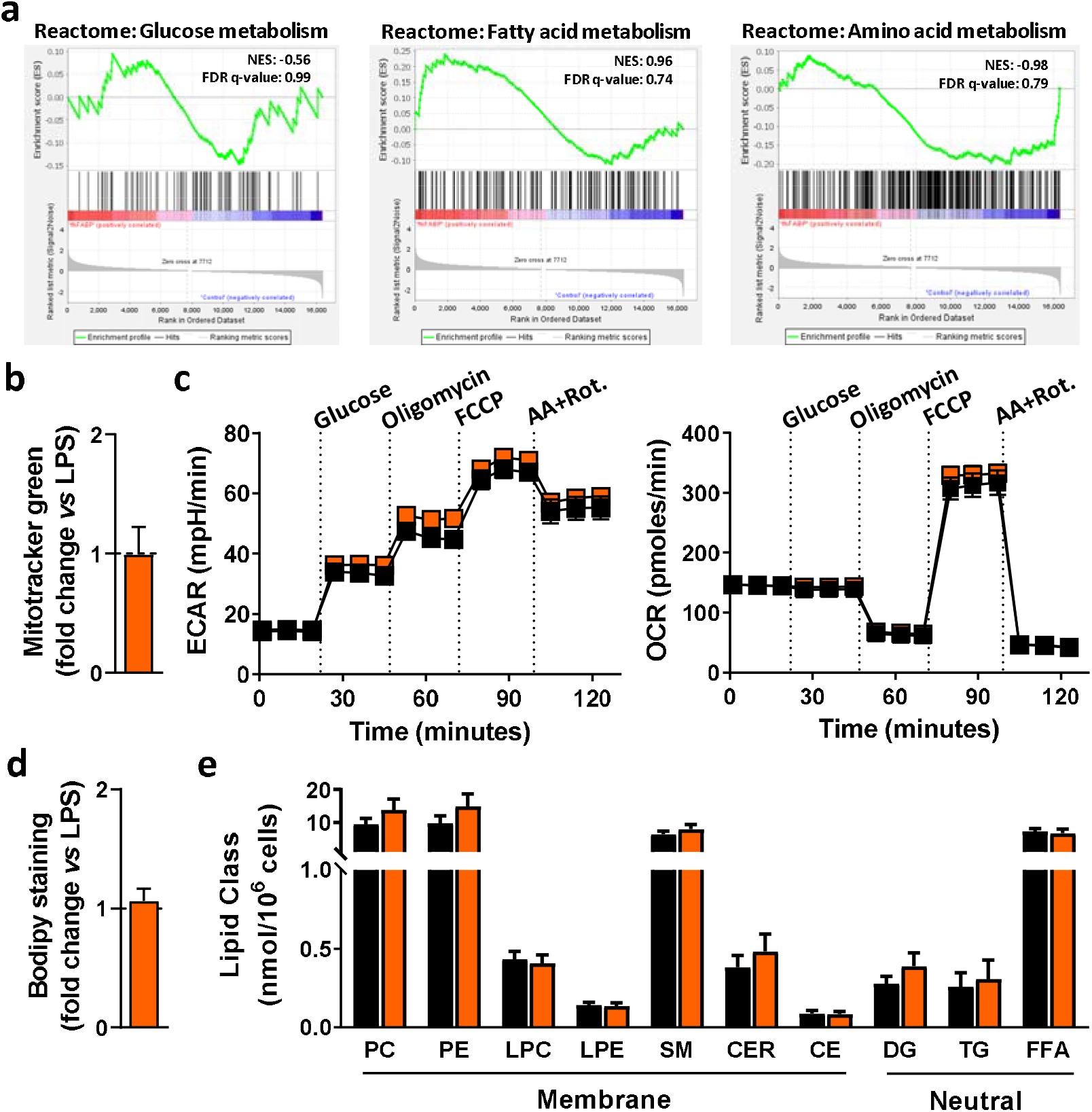
*Fasciola hepatica* FABP1 does not affect intrinsic metabolism of dendritic cells. Human moDCs were stimulated for 5 h with PBS (black squares/bars) or 25 μg/ml recombinant *fh*FABP1 (orange squares/bars) in presence of 100 ng/ml LPS. Gene set enrichment analyses were done on the RNAseq data from Figure 5 using the Reactome module for the main metabolic pathways **(a).** The intrinsic metabolic phenotype of moDCs were determined by measuring extracellular acidification rates (ECAR) and oxygen consumption rates (OCR) with a Seahorse flux analyser **(c).** The mitochondrial mass **(b)** and intracellular neutral lipid content **(d)** were determined by flow cytometry, using Mitotracker green and Bodipy staining, respectively. The lipid class concentrations were also quantified by targeted lipidomics using the Lipidyzer platform **(e).** PC, Phosphatidylcholine; PE, Phosphatidylethanolamine; LPC, Lysophosphatidylcholine; LPE, Lysophosphatidylethanolamine; SM, Sphingomyelin; CER, Ceramides; CE, Cholesterylester; DG, Diglycerides; TG, Triglycerides; FFA, Free-fatty acids. Results are expressed as mean ± SEM (n=3-5 independent experiments).

**Figure S7.**
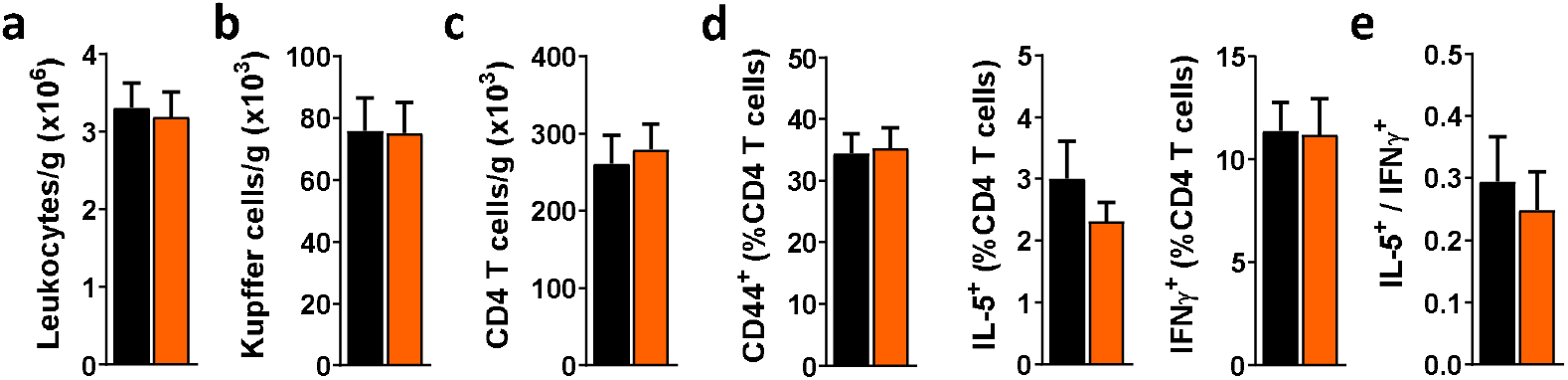
*Fasciola hepatica* FABP1 does not induce type 2 immune response in livers from obese mice. Mice were fed a high-fat diet (HFD) for a period of 12 weeks and next received intraperitoneal injections of 25 μg recombinant *fh*FABP1 (orange squares/bars) or vehicle (PBS, black squares/bars) every 3 days for 4 weeks, as described in Figure 8. The number of liver total leukocytes **(a),** Kupffer cells **(b)** and CD4^+^ T cells **(c),** together with the percentage of CD44^+^, IL-5^+^, IFNγ^+^ CD4^+^ T cells **(d)** were determined by flow cytometry. The IL-5^+^-on-IFNγ^+^ ratio was calculated **(e).** Results are expressed as mean ± SEM. # P ≤ 0.05 *vs* HFD (n=5-6 mice per group).

